# Predicting cell-type-specific non-coding RNA transcription from genome sequence

**DOI:** 10.1101/2020.03.29.011205

**Authors:** Masaru Koido, Chung-Chau Hon, Satoshi Koyama, Hideya Kawaji, Yasuhiro Murakawa, Kazuyoshi Ishigaki, Kaoru Ito, Jun Sese, Yoichiro Kamatani, Piero Carninci, Chikashi Terao

## Abstract

Transcription is regulated through complex mechanisms involving non-coding RNAs (ncRNAs). However, because transcription of ncRNAs, especially enhancer RNAs, is often low and cell type-specific, its dependency on genotype remains largely unexplored. Here, we developed <u>m</u>utation <u>e</u>ffect prediction on <u>n</u>cRNA <u>tr</u>anscription (MENTR), a quantitative machine learning framework reliably connecting genetic associations with expression of ncRNAs, resolved to the level of cell type. MENTR-predicted mutation effects on ncRNA transcription were concordant with estimates from previous genetic studies in a cell type-dependent manner. We inferred reliable causal variants from 41,223 GWAS variants, and proposed 7,775 enhancers and 3,548 long-ncRNAs as complex trait-associated ncRNAs in 348 major human primary cells and tissues, including plausible enhancer-mediated functional alterations in single-variant resolution in Crohn’s disease. In summary, we present new resources for discovering causal variants, the biological mechanisms driving complex traits, and the sequence-dependency of ncRNA regulation in relevant cell types.

## INTRODUCTION

Large scale human transcriptome analyses have revealed that long-ncRNA (lncRNA) and enhancer RNA expression is tissue- and cell-type-specific and regulates human development and homeostasis ((DGT) et al., 2014; Andersson et al., 2014; Hon et al., 2017). Genome-wide association studies (GWAS) have found many complex traits-associated variants enriched in loci from which ncRNAs are transcribed (Andersson et al., 2014; Boyd et al., 2018; Hon et al., 2017; Kristjánsdóttir et al., 2018; Maurano et al., 2012) as well as substantial heritability enrichments in transcribed enhancer regions (Finucane et al., 2015). These findings implicate ncRNAs in the mechanisms driving complex traits. GWAS has allowed inference of trait-relevant tissues or cell-types (Finucane et al., 2015, 2018), biological pathways (Iotchkova et al., 2019; Lamparter et al., 2016), and therapeutic drugs (Terao et al., 2016); therefore, determining the influence of non-coding causal variants from GWAS on ncRNA expression in relevant cell-types is a promising approach to deepen our understanding of complex trait mechanisms (Figure 1A). For mRNAs, such analyses have already improved understanding of the genetic architecture of complex traits; mRNA expression quantitative trait loci (eQTL) information, resolved to the tissue- or cell-type-level, has been linked to GWAS results to implicate the cell types and tissues involved in complex traits (Ardlie et al., 2015; Ishigaki et al., 2017). However, mapping eQTL in a given tissue or cell-type has traditionally required gene expression datasets for the target tissue or cell-type, as well as genotypes from, in general, over 100 individuals (Figure S1A). Discovery of eQTL for ncRNAs is especially challenging due to their often low expression levels and high cell-type-specificity (Andersson et al., 2014; Hirabayashi et al., 2019; Hon et al., 2017).

**Figure 1.**
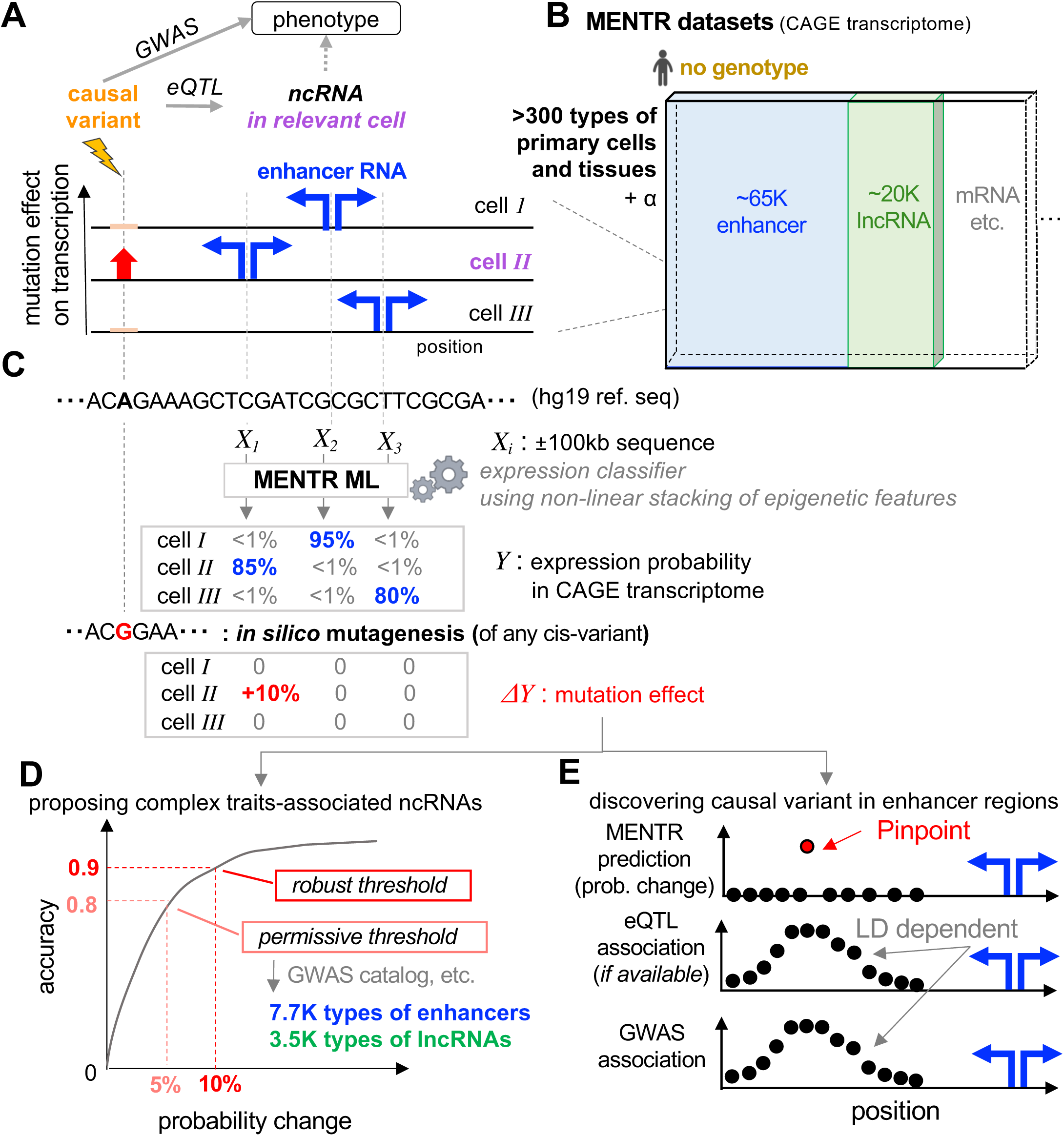
Strategy to find variants’ effects on ncRNA transcription by MENTR. (A) MENTR accurately predict effects of any mutation on ncRNA transcription in a cell-type dependent manner, enabling interpretation of complex trait-associated loci. Active enhancers, bidirectionally transcribed in a cell-type dependent manner, were shown as representative example of ncRNA. (B) Available transcriptome datasets for training MENTR ML models. CAGE transcriptome datasets other than those collected by FANTOM5 can be used. (C) MENTR ML models learn promoter and enhancer-level on-off patterns from the TSS ± 100-kb genome sequence. After the training ML models, mutation effects of any mutations on proximal promoters and enhancers can be inferred by comparing predicted expression probability from different input genome sequences (*in silico* mutagenesis). (D) Robust and permissive MENTR predictions can be selected by the threshold determined by comparison with previous genetic studies. Thousands of complex traits-associated ncRNAs were catalogued by using previous GWAS findings. (E) Schematic of pinpoint mutation effects by MENTR, compared with GWAS findings and conventional eQTL findings.

Recently, machine learning (ML) models, including convolutional neural networks (CNN), have succeeded in predicting epigenetic events (Hoffman et al., 2019; Zhou and Troyanskaya, 2015) and gene expression levels in a tissue- or cell-type-dependent manner (Kelley et al., 2018; Zhou et al., 2018). This advance has been made by extending conventional, short-range motif analysis and performing ML on kilobase (kb) scale genome sequence patterns. Despite training ML models without individual genotype data (in other words, models assume that the training data comes from cells with the reference genome sequence), *in silico* mutagenesis can predict the effect of genetic variants on a particular transcript. Estimates of mutation effect made based on ML have been comparable with those based on QTL studies for mRNA expression (Kelley et al., 2018; Zhou et al., 2018). Notably, *in silico* mutagenesis has achieved comparable performance despite not requiring genotype data, thus accumulated transcriptome datasets may be suitable for training. Furthermore, *in silico* predictions are not affected by linkage disequilibrium (LD), allowing pin-point prediction of causal variants on mRNA transcriptional changes (Zhou et al., 2018). These previous studies motivated us to expand the ML-based framework to cell-type-specific expression of ncRNA (Figure 1A, Figure S2A). Here we demonstrate MENTR (<u>m</u>utation <u>e</u>ffect prediction on <u>n</u>cRNA <u>tr</u>anscription), a ML program trained using cell-type-specific ncRNA and enhancer transcription, measured by cap analysis of gene expression (CAGE), that can accurately predict the effect of mutations on ncRNA expression ((DGT) et al., 2014; Andersson et al., 2014).

## RESULTS

### Strategy to predict mutations’ effects on ncRNAs

MENTR ML models learn cell-type-specific transcription of promoters (including >20K types of lncRNA) and >65K types of enhancers from only +/- 100-kb human reference genome sequence (hg19) surrounding transcriptional start site (TSS) (Figure 1B and 1C). We developed MENTR ML models by combining deep convolutional neural networks (from 2-kb sequence bin to 2002 epigenetic features, using publicly available pre-trained models (Zhou and Troyanskaya, 2015; Zhou et al., 2018)) and binary classifiers using non-linear, gradient boosting trees (from the many epigenetic features in +/- 100-kb sequence to accurate transcription probability; see METHOD DETAILS) (Chen and Guestrin, 2016). The binary classifier outputs a probability of expression for each tissue or cell-type, chosen because we focus here on predicting lowly-expressed RNAs whose quantitative measurement might be not reliable (Hirabayashi et al., 2019)). We trained the MENTR ML models using the autosomal mRNA and ncRNA promoter- and enhancer-level transcripts profiled by CAGE except for chromosome 8 and tested the accuracy using those of chromosome 8 (Figure S1B). After training, MENTR can predict mutation effects on transcripts (for each tissue or cell-type used for training) by *in silico* mutagenesis, the estimates of which are prioritizable based on the degree of probability change (Figure 1D) and are free from LD structure (Figure 1E).

### Accurate prediction of cell-type-specific promoter- and enhancer-level transcription

We trained MENTR ML models using FANTOM5 CAGE transcriptome data from 347 types of samples comprising a variety of primary cells and tissues (see METHOD DETAILS). Training MENTR using +/- 100-kb flanking the transcriptional start site (TSS) achieved the highest accuracy, and shorter input sequences slightly decreased accuracy (Figure S3). Interestingly, training based on only promoter expression could predict enhancer RNA expression, and vice versa, supporting that promoters and enhancers at least partially share sequence-based regulatory machineries. Training using only enhancer data slightly decreased predictive accuracy for both promoters and enhancers, presumably due to technical limitation of accurate detection of enhancer RNAs. Thus, we decided to train using both promoters and enhancers, thereby including RNAs expressed over a wide range of abundances. To reduce computational costs, we used linear penalized logistic regression models using boosting (MENTR_linear_) (Bühlmann, 2006; Chen and Guestrin, 2016) as a binary classifier in the above screening procedures.

MENTR revealed good prediction accuracy for enhancer expression in a probability-dependent manner (Figure 2A). The area under the receiver operating characteristic curve (AUROC) was 0.69 ± 0.05 for enhancers in 347 sample ontologies, 0.76 ± 0.04 for lncRNAs. Unsurprisingly, AUROC values for coding mRNA (0.83 ± 0.02) were higher than those for ncRNA, and predictions of small RNA, pseudogene, and short ncRNA expression were not always accurate (Figure 2B, Table S1, and Table S2). Even though MENTR was trained using expression as an on-off variable, the correlation between predicted expression probability and measured expression levels in FANTOM5 CAGE transcriptome datasets was much higher than the previous method ExPecto (Zhou et al., 2018) for lncRNA (Wilcoxon signed rank test P = 1.8×10^-58^) and mRNA (P = 1.3×10^-58^) (Figure S2B). ExPecto uses transcription strand information and thus cannot be used to predict bidirectionally-transcribed enhancer RNAs (see METHOD DETAILS). These data indicated that MENTR was suitable for accurate prediction of lowly-expressed enhancers and lncRNAs, not only mRNAs, in CAGE transcriptomes.

**Figure 2.**
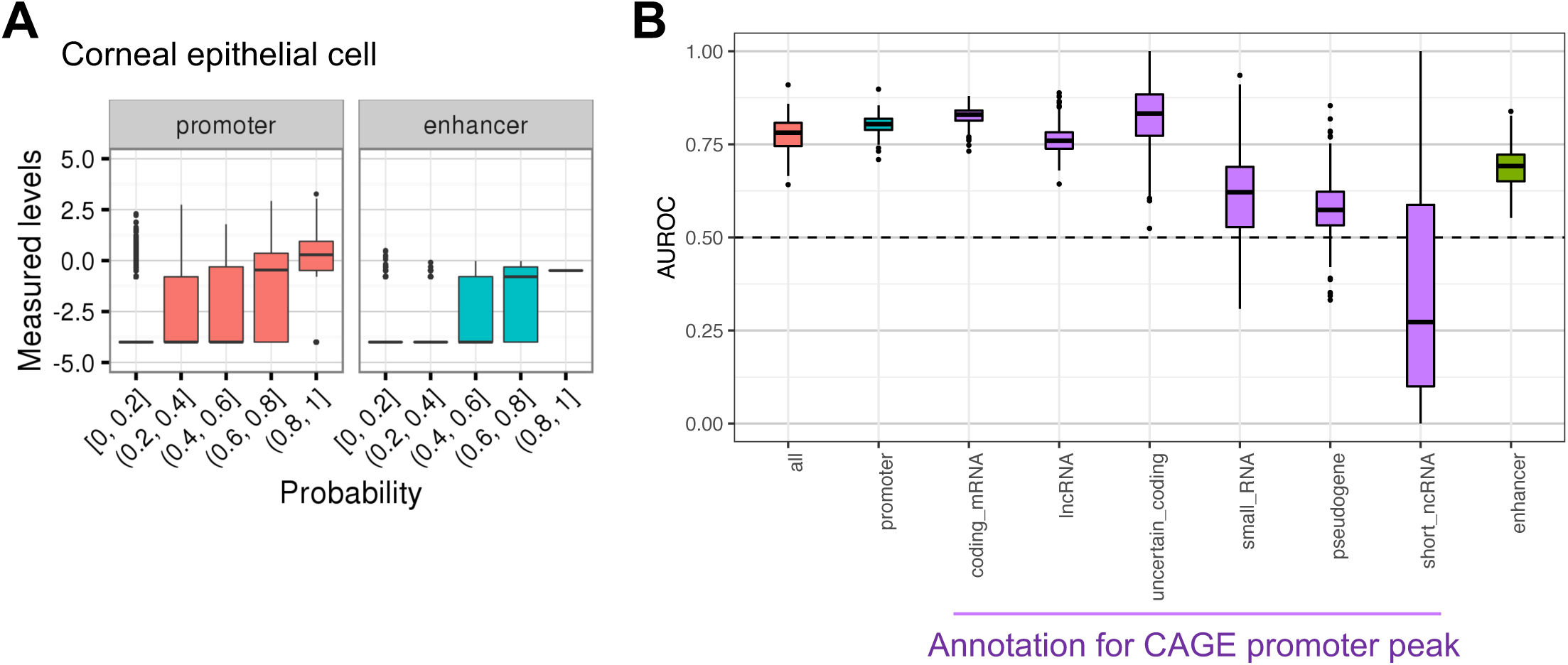
Cell-type-specific prediction of promoter- and enhancer-level expression by MENTR. (A) Distribution of measured expression levels for the bins of probability from the trained MENTR ML model trained by CAGE transcriptome of corneal epithelial cell (a representative result). (B) Summary of predictive accuracy (AUROC) for the FANTOM5 347 sample ontologies. Purple boxes show the AUROC for the only promoters including the annotation indicated in x-axis. The box plots show the first and third quartiles, the center line represented the median, the upper whisker extended from the hinge to the highest value that is within 1.5 × IQR (inter-quartile range) of the hinge, the lower whisker extended from the hinge to the lowest value within 1.5 × IQR of the hinge, and the data beyond the end of the whiskers were plotted as points. Random accuracy (0.5) was shown as a dashed line.

**Table 1.** GWAS traits-associated promoters and enhancers. Threshold *None*, all of the tested data; Threshold *Permissive*, analysis results whose absolute *in silico* mutation effects were ≥ 0.05; Threshold *Robust*, analysis results whose absolute *in silico* mutation effects were ≥ 0.1; #Tested pairs, the number of combination of variant, trait, CAGE peak, and MENTR ML models; #variants, the distinct number of tested GWAS variants and variants in LD; #CAGE peaks, the number of CAGE promoters and enhancers.

| Threshold | #Tested pairs | #variants | #CAGE peaks |  |  |  |  |
| --- | --- | --- | --- | --- | --- | --- | --- |
|  |  |  | Total |  |  |  |  |
|  |  |  |  | promoter | mRNA | lncRNA | enhancer |
| *None* | 433,072,776 | 41,223 | 177,734 | 131,982 | 542,984 | 85,480 | 45,752 |
| *Permissive* | 197,306 | 17,306 | 32,979 | 25,204 | 24,502 | 3,548 | 7,775 |
| *Robust* | 15,395 | 1,256 | 2,067 | 1,693 | 1,235 | 168 | 374 |

Interestingly, prediction of transcription of promoters annotated as “CpG-less” was accurate (AUROC: 0.73 ± 0.02) but underperformed those annotated as containing CpGs (Wilcoxon-Mann-Whitney test P = 2.3×10^-178^) (Figure S4). This suggests that MENTR ML models have learned the importance of CpG sites for transcription without any prior knowledge, but also learned the exception rules (transcription from CpG-less sites) at the same time. Moreover, transcription of promoters annotated as “TATA-less” were rather more predictable than those annotated as including TATA boxes (Figure S4; P = 2.2×10^-31^). Recent studies have shown the existence of many TATA-less promoters in mammals, with transcription initiating in a different and more predictably flanking-sequence-dependent manner compared to TATA-containing promoters (Anish et al., 2009; Donczew and Hahn, 2017). Taken together, MENTR could learn genome sequence patterns that influence transcription. Therefore, interpreting the output of MENTR might be useful for understanding the biology of gene expression.

Considering the quite low expression levels of many ncRNAs, we were concerned that MENTR could learn anomalous read patterns, noise, or artifacts related to transcript mappability that vary depending on sequence context, rather than reproducible transcriptional status; if this were the case, it would be unexpected if MENTR could accurately predict the expression patterns of RNAs that are in fact transcribed, but are not detected due to low depth of sequencing or degradation. To evaluate this possibility, we took advantage of enhancer RNA transcription data in five ENCODE cell lines profiled using NET-CAGE, a sensitive method for measuring nascent RNA transcription (Hirabayashi et al., 2019). We then evaluated false positive (FP) predictions, in which a transcript was not detected by the standard CAGE method, but MENTR predicted a probability of expression >0.5 (based on ML models trained using standard CAGE data). We observed that 31-70% of the transcripts which were initially considered as FP predictions were actually transcribed across all five cell lines when assayed using NET-CAGE (Table S3; P < 3.6×10^-3^). These results indicate that MENTR ML models accurately learned sequence-dependent transcription patterns for enhancers, even though a substantial fraction of them might have been undetectable in the standard CAGE transcriptomes used for training. This indicates that we might underestimate the accuracy of MENTR, especially for enhancers (and possibly small RNAs, pseudogenes, and short ncRNA) by AUROC values (Figure 2, Table S1, and Table S2) using standard CAGE data to define ground truth. Thus, we did not filter out MENTR ML models by their AUROC values in the following analyses.

### Verification of predicted mutation effects by comparing with previous eQTL/caQTL studies

Having trained MENTR to predict RNA transcription based on genomic sequence, we next sought to evaluate its ability to predict the effect of mutations on transcription. In order to verify the accuracy of predictions based on *in silico* mutagenesis, we additionally trained a MENTR ML model on CAGE transcriptomes of lymphoblastoid cell lines (LCL) (AUROC for promoter = 0.82 and that for enhancer = 0.71; Table S4) for which eQTL, as well as chromatin accessibility QTL (caQTL), were available (Garieri et al., 2017; Kumasaka et al., 2019). We compared *in silico* mutation effects predicted by MENTR and the effect sizes in the two previous QTL studies for the alternative alleles.

First, we compared *in silico* mutation effects from MENTR ML models trained on the 347 FANTOM5 CAGE transcriptomes with eQTL coefficients in LCL on promoter- and enhancer-level CAGE transcriptome analysis (Garieri et al., 2017) (Figure S5A). We found that the higher the predicted mutation effects by the MENTR ML model trained using LCL data, the more consistent with the observed effect of the variant in the eQTL study. This was true even for enhancers and lncRNAs (Figure 3A-C and Figure S6A; see red line). While the mutation effects predicted in the LCL model did not always provide the best accuracy, the models giving good accuracy were trained on transcriptional profiles highly correlated with that of LCL, including lymphocyte in FANTOM5 (Figure 3A-C and Figure S6A; see orange lines in heatmap colors). Prediction accuracy among the different models were variable, especially for low abundance transcripts. Bad prediction in models trained using cells or tissues with transcription profiles dissimilar to LCLs was obvious for enhancer and lncRNA (Figure 3A-C; see blue lines in heatmap colors). These results indicated that cell-type-specific training is important for accurate prediction of mutation effects, especially for enhancer and lncRNA.

**Figure 3.**
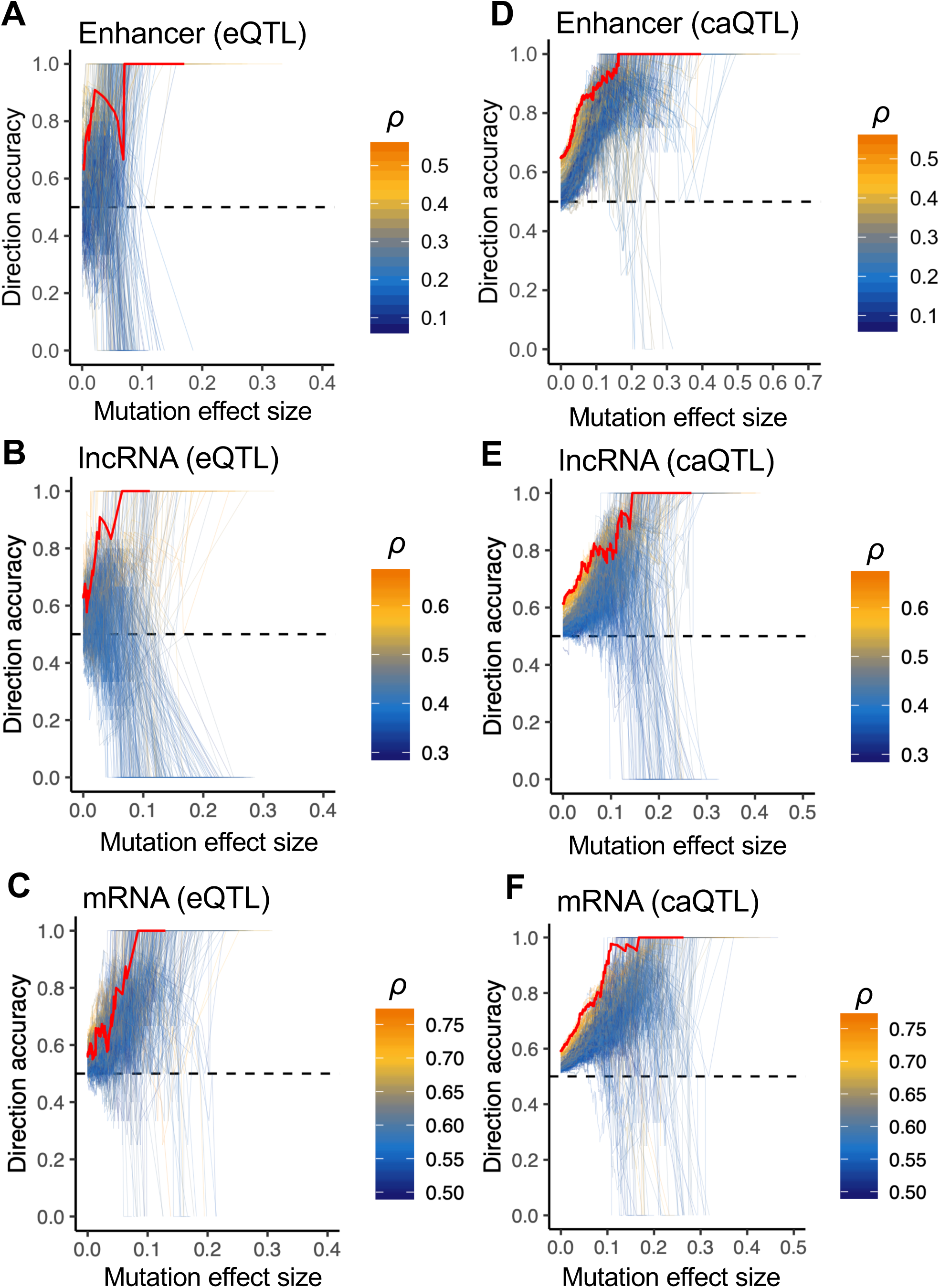
Accurate MENTR predictions of mutation effects on ncRNA expression in a cell-type-dependent manner. Concordance rate (y-axis) of directions between *in silico* mutation effect (*β_mutgen_*) and effect size from the QTL study (*β_QTL_*) at the threshold of absolute *β_mutgen_* (x-axis). *β_QTL_* was obtained from LCL CAGE QTL study (Garieri et al., 2017) in (A–C) and from LCL caQTL study (Kumasaka et al., 2019) in (D–F). Accuracy for enhancers was shown in (A, D), that for lncRNA promoters was in (B, E), and that for mRNA promoters was in (C, F). These *β_QTL_* values were compared with *β_mutgen_* values from the MENTR ML model trained by LCL CAGE transcriptome (red line) as well as models trained by FANTOM5 347 CAGE transcriptomes (heatmap color, indicating transcriptomic correlation(Spearman’s ρ) with LCL CAGE transcriptome). The bluer, the more unmatched comparison of cell-types. Random accuracy (0.5) was shown as a dashed line.

Second, we compared *in silico* mutation effects with coefficients of caQTL (Kumasaka et al., 2019) under the hypothesis that transcription of CAGE promoter- and enhancer-regions were regulated similarly to chromatin accessibility, as measured by ATAC-seq peaks (Figure S5B). Testing on the variant-promoter and variant-enhancer pairs supports the same conclusion as the analysis using eQTL (Figure 3D-F, Figure S6B), indicating that prediction accuracy of MENTR is not dependent on the CAGE method and that chromatin accessibility is regulated in a sequence-dependent manner similar to that of transcription of enhancers and promotors. Although our predictions were based on binary transcription state, the *in silico* mutation effect sizes were correlated with the effect size in the caQTL study for both promoters (Spearman’s ρ = 0.25 [95% CI: 0.24-0.26]) and enhancers (Spearman’s ρ = 0.36 [95% CI: 0.32-0.36]) (Figure S7). Taken together, this shows that MENTR accurately predicts transcription, including enhancers, in a cell-type-dependent manner, especially when we used predictions with higher mutation effect size. Hereafter, we defined the permissive threshold achieving ∼80% concordance and the robust threshold achieving >90% concordance in Figure 3A-C as 0.05 and 0.1 of absolute *in silico* mutation effects, respectively.

Next, we analyzed 26 types of tissues assayed by CAGE in FANTOM5 and also in GTEx (manually matched; Table S5) and compared results between *in silico* mutagenesis by MENTR and GTEx eQTL. We limited this analysis to variants within 1-kb of transcripts, because almost all non-zero mutation effects were obtained +/- 1-kb from TSS (for example, 19,376 out of 86,724 variant-promoter pairs showed non-zero mutation effects, from which 18,226 pairs (94.0%) were in +/- 1-kb from TSS (Table S6)). We aggregated the promoter-level predictions of mutation effects into gene-level predictions (Figure S8A) and, by comparing with GTEx eQTL estimates, verified that MENTR’s prediction, at both permissive and robust thresholds, were accurate in almost all types of tissues (Figure S8B; Wilcoxon signed rank test P = 9.8×10^-7^ for permissive threshold and P = 4.2×10^-7^ for robust threshold). These results strongly support the applicability of *in silico* mutagenesis using MENTR to predict the effect of genetic variants on transcription in many types of tissues.

Seeking additional evidence to support the predictive validity of MENTR, we analyzed variants whose effects on mRNA transcription had been tested using reporter assays (Zhou et al., 2018). We predicted *in silico* mutation effects of the rs147398495 (chr3:46249943 GTTC>G; *CCR1* locus) and rs381218 (chr6:32977420:G>T; *HLA-DOA* locus) and the variants in LD (r^2^ ≥ 0.2 in 1KGp3v5 EUR) on the representative promoter. Interestingly, the variants with the strongest *in silico* mutation effects were the lead variants themselves (rs147398495 and rs381218) as supported by experimental evidence from the previous study (Zhou et al., 2018). Importantly these were also the only variants satisfying the robust threshold of *in silico* mutation effect (Figure S9).

### Cataloging and prioritizing GWAS findings by MENTR

We evaluated *in silico* mutation effects of previously reported GWAS variants (in GWAS catalog and our previous studies; Akiyama et al., 2017; Ishigaki et al., 2019; Kanai et al., 2018) as well as variants in LD with the variants (r^2^ ≥0.7, see METHOD DETAILS) to facilitate interpretation of GWAS results that may alter ncRNA transcription. As a result, we identified over ten thousand (permissive threshold) or ∼500 (robust threshold) ncRNAs associated with GWAS traits (Table1 and Table S7-S9). We *released* all these results (GWAS trait-associated ncRNA database) as publicly available resources in a user-friendly GUI application for both Windows and Mac users (*url*). (will do *after acceptance*)

Among 42 complex diseases catalogued in Biobank Japan, 36 diseases showed positive correlations between effect sizes of risk variants and their *in silico* mutation effects (after conditioning on MAF, transcription type (promoter or enhancer) and distance between variants and transcripts). Among them, 15 diseases, including rheumatoid arthritis (RA), Graves’ disease (GD), chronic hepatitis B (CHB), and pancreatic cancer (PaCa), showed significant correlations (Bonferroni-corrected level of significance (P < 0.05/42)) (Figure 4 and Table S10). This indicates alteration of ncRNA expression by the variants that underlies risk for these complex diseases.

**Figure 4.**
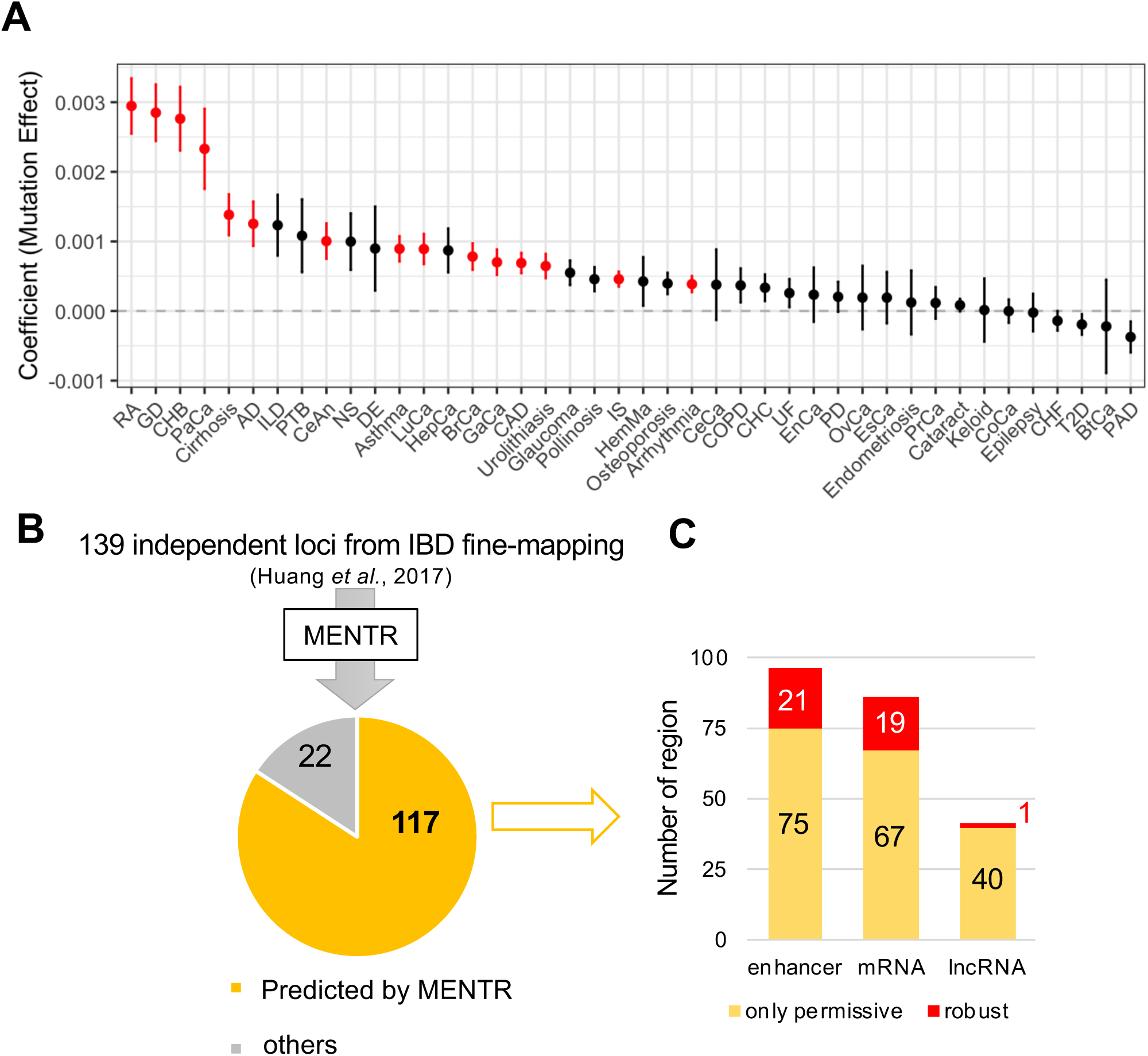
MENTR predictions to explain and prioritize GWAS findings. (A) Association of *in silico* mutation effect sizes with GWAS effect sizes. Regression coefficients of mutation effects on GWAS effect size for the indicated complex diseases (Ishigaki et al., 2019) in x-axis. The coefficients were conditioned on MAF of variants in testing samples (East Asian), absolute variant-TSS distance, and dummy variable for promoter or enhancer. The statistically significant results (Bonferroni-corrected level of significance; P < 0.05/42) were highlighted (red). The error bars show standard error of regression coefficient. (B) The number of credible loci of IBD (Huang et al., 2017), whose causal variants were predicted as ‘permissive’ by MENTR in at least one cell-type or tissue. (C) The number of credible loci, predicted by MENTR for the indicated type of transcription. AD, Atopic dermatitis; BrCa, Breast cancer; BtCa, Biliary tract cancer; CAD, Coronary artery disease; CeAn, Cerebral aneurysm; CeCa, Cervical cancer; CHB, Chronic hepatitis B; CHC, Chronic hepatitis C; CHF, Congestive heart failure; CoCa, Colorectal cancer; COPD, Chronic obstructive pulmonary disease; DE, Drug eruption; EnCa, Endometrial cancer; EsCa, Esophageal cancer; GaCa, Gastric cancer; GD, Graves’ disease; HemMa, Hematological malignancy; HepCa, Hepatocellular carcinoma; ILD, Interstitial lung disease; IS, Ischemic stroke; LuCa, Lung cancer; NS, Nephrotic syndrome; OvCa, Ovarian cancer; PaCa, Pancreatic cancer; PAD, Peripheral artery disease; PD, Periodontal disease; PrCa, Prostate cancer; PTB, Pulmonary tuberculosis; RA, Rheumatoid arthritis; T2D, Type 2 diabetes; UF, Uterine fibroids.

We further found that the MENTR could suggest plausible functional roles of GWAS variants for complex traits. We evaluated mutation effects of 95% credible sets of over 4000 causal variants in 139 independent loci from the fine-mapping study on inflammatory bowel disease (IBD) (Huang et al., 2017). Surprisingly, we found that the credible sets in 117 loci (84%) included variants with an effect on transcription predicted by *in silico* mutagenesis by MENTR (Figure 4B). In contrast, MENTR generally predicted no effect for >70% of non-eQTL variant-transcript pairs near GTEx eQTL (Table S6). In these predictions, effects on transcription of enhancer and mRNA explained 96 and 86 loci, respectively (Figure 4C), reaffirming the importance of transcribed enhancers in the etiology of complex traits.

Among the robust predictions of IBD credible sets (Table S11), MENTR pinpointed the best candidate of causal variants whose effects on transcriptional activity were experimentally validated in previous studies. In one example, rs17293632 in the *SMAD3* region showed the highest posterior probability (40%) in fine-mapping study of Crohn’s disease (CD) and is also known to be associated with risk of asthma and coronary artery disease (CAD) (Demenais et al., 2018; Huang et al., 2017; Turner et al., 2016) (Figure 5A), providing interpretation of the association in the context of cell-type-specific ncRNA transcription profiles. There are many variants showing strong eQTL effects on *SMAD3* near rs17293632 (Figure 5B) making it difficult to identify a causal variant based on eQTL associations, and the LD structure of this region further complicates biological interpretation. On the other hand, MENTR predicted that the T allele of rs17293632 (a risk allele for Crohn’s disease and asthma, and a protective allele for CAD) decreased expression of the enhancer ADDG15067442347.E, in many types of cells plausibly relevant to these diseases: colon, neutrophil, natural killer cell, eosinophil, and macrophage (Fahy, 2009; Wéra et al., 2016; Yadav et al., 2011) (Figure 5C and Table S12). Consistent with this prediction, rs17293632 is located at an active enhancer region (a H3K27ac ChIP-seq peak) of many types of cells including human sigmoid colon, rectal mucosa, monocytes, coronary artery smooth muscle cells. The T allele is known to lower chromatin accessibility and decrease binding of AP-1 transcription factor (Farh et al., 2015; Huang et al., 2017; Miller et al., 2016; Turner et al., 2016), supporting enhancer-mediated transcriptional regulation by this variant. Notably, MENTR predicted that rs17293632 and variants in LD (r^2^ ≥0.2 in 1KGp3v5 EUR) would have no direct effects on transcription from *SMAD3’s* representative promoter (Figure 5C). rs17293632 was predicted to affect transcription from three minor promoters (p6, p11, p13@SMAD3) in corneal epithelial cell and CD14^+^CD16^+^ monocyte (Table S11), suggesting that the alternative promoter usage might be regulated by the variant.

**Figure 5.**
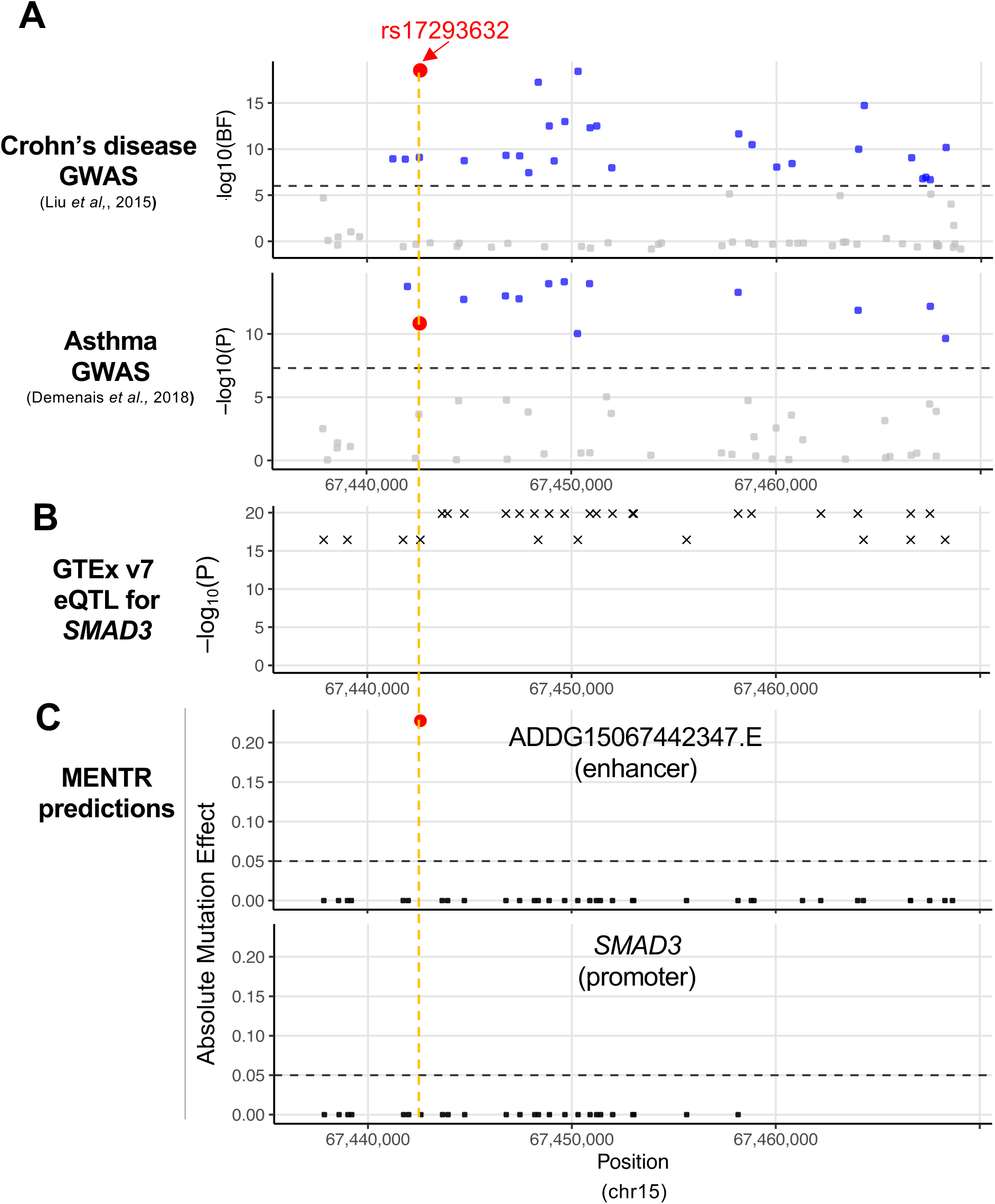
MENTR illuminating a candidate of a causal variant in Crohn’s disease by linking variants with transcribed enhancers in relevant cell-types. (A) Association plots from Crohn’s disease transethnic meta GWAS (Liu et al., 2015) and asthma transethnic meta GWAS results (Demenais et al., 2018). Bayes Factor (BF) was used in the former, and p-value was used in the latter. BF threshold in the paper (log_10_BF = 6) and Genome-wide significant threshold (P = 5×10^-8^) were shown as a dashed line, respectively. (B) Association plot for *SMAD3* eQTL from GTEx v7 eQTL studies. Significant eQTL determined by GTEx (qval < 0.05) was shown. The minimum p-value for each locus over tissues was used. (C) *in silico* mutation effects for the indicated transcripts from MENTR ML model for neutrophil. Absolute mutation effect sizes were shown. Permissive threshold of mutation effect (0.05) was shown as a dashed line. *SMAD3*, representative promoter (p1@SMAD3).

Similarly, we found two additional plausible examples from the IBD credible sets. First, MENTR predicted that rs713875 (near *LIF* and *HORMAD2* loci) lowered the expression of enhancer ADDG22030591785.E in CD-relevant CD8^+^ - T cell (Kadivar et al., 2016) α - β (Figure S10 and Table S11), whose alternative allele was methylated in whole blood (Hutchinson et al., 2014). Second, MENTR predicted that the alternative allele of rs4456788 (near *ICOSLG* loci) lowered the expression of enhancer ADDG21045616099.E in CD-relevant cell types such as CD4^+^ - T cell (Imam et al., α - β 2018) (Figure S10 and Table S11), whose alternative allele decreased reporter activity of surrounding sequence in a cell line model (Mattioli et al., 2019). MENTR can thus help prioritize plausible causative variants evidenced by previous findings and raise meaningful new hypotheses based on GWAS results.

## DISCUSSION

We developed MENTR to predict the effect of genetic variants on transcription, including transcription of ncRNAs. We demonstrated that *in silico* mutations predicted to have strong effects were highly concordant with the observed effects of known variants in a cell-type-dependent manner. As an example of how MENTR can facilitate interpretation of GWAS results and raise new mechanistic hypotheses, we pinpoint variants previously short-listed as credible causative variants and suggest plausible enhancer-mediated functional interpretations. In addition to the pre-calculated mutation effects of GWAS variants in CAGE transcriptomes from >300 types of human primary cells and tissues, we release programs which will allow others to predict the effect of any variant, in any region of interest, on ncRNA as well as mRNA transcription (*will available after acceptance*).

An important feature of MENTR compared with previous relevant ML methods is the successful expansion of *in silico* mutagenesis principle into transcribed ncRNA, especially enhancers, which are bidirectionally transcribed from active enhancer regions and enrich GWAS variants in a pathologically relevant cell types (Andersson et al., 2014; Kim et al., 2010; Murakawa et al., 2016). So far, 65K transcribed enhancers are known; but their expression has not been used for training ML models (Kelley et al., 2018; Zhou et al., 2018) despite their known high cell-specificity and trait relevance (Andersson et al., 2014; Hon et al., 2017). MENTR was trained using binarized RNA transcription patterns, yet surprisingly our approach provided much more quantitative predictions than the previous method which learned expression levels along a quantitative spectrum (Figure S2).

Recently, the clinical importance of enhancers have been widely recognized, as shown in IBD (Boyd et al., 2018) and cancer studies (Zhang et al., 2019). Nevertheless, the functional roles of variants on or near enhancers are unclear, and only a few enhancer RNA QTL studies in LCL using limited number of subjects (154 European (Garieri et al., 2017) and 69 Yoruban (Kristjánsdóttir et al., 2018)) have been conducted. Although enhancer RNA QTL might be partly discovered from ChIP-seq for histone marks (*e.g.*, H3K4me1 and H3K27ac), DNase-seq or ATAC-seq, these studies have also been conducted for LCL, blood cells, and other limited types of cells using only hundreds of European or Yoruba individuals (Alasoo et al., 2018; Banovich et al., 2018; Bryois et al., 2018; Chen et al., 2016; Degner et al., 2012; Delaneau et al., 2019; Gate et al., 2018; Kumasaka et al., 2019; Pelikan et al., 2018). An alternative approach is the experimental identification of transcribed enhancers (van Arensbergen et al., 2019), but it would still be challenging to conduct such experiments in many types of cells. Thus, the present study provides the only resource for interpreting the genetic regulation of enhancer transcription in >300 types of primary cells and tissues.

In GTEx, currently the largest eQTL database, the number of donors is at most 500, depends heavily on tissue accessibility, and over 80% of donors are European (GTEx v7) (Aguet et al., 2017). Therefore, the effects of low-frequency or rare variants remain unknown, along with population-specific variants in non-Europeans.

Furthermore, eQTL strongly depends on LD structure in the tested population. On the other hand, ML-based *in silico* mutagenesis trained using the human reference genome sequence is free from allele frequency and LD issues and instead prioritizes causal variants at single basepair resolution (Zhou et al., 2018). MENTR-predicted mutation effects on transcription are zero in many cases, but provided pinpoint estimation of causal variants without depending on LD structure (Figure 4-5, and Table S6). MENTR is thus a useful new tool to filter GWAS variants in LD to identify the most likely causal variants.

Planned large scale whole-genome sequencing projects and increasing GWAS samples sizes (Saunders et al., 2019) are expected to increasingly reveal infrequent and rare variants associated with complex traits. These variants will increasingly not be contained in eQTL catalogs thus other tools are needed to filter credible variants and make testable hypotheses about the mechanisms by which these variants drive disease, or other phenotypes of interest. MENTR is an attractive first choice to provide clues and interpretations of such non-coding GWAS associations, especially for low-frequency variants.

## Supporting information

Key Resources Table

## Acknowledgments

We deeply thank Dr. Nicholas Parrish for critically reviewing and editing the manuscript. We thank FANTOM consortium members for providing datasets and valuable discussions. Computational resource of AI Bridging Cloud Infrastructure (ABCI) provided by the National Institute of Advanced Industrial Science and Technology (AIST) was used for *in silico* mutagenesis.

## Author Contributions

M.K, C-C.H, Y.K and C.T conceived the study. M.K conducted analysis with the help of C-C.H, S.K, K.Ishigaki, K. Ito and C.P. C-C.H analyzed CAGE transcriptome data. H.K and Y.M analyzed NET-CAGE transcriptome data. M.K and C.T wrote the manuscript. J.S contributed to providing GPU computational resources which were necessary for the present study. P.C and C.T supervised this study.

## STAR Methods

### LEAD CONTACT AND MATERIALS AVAILABILITY

Further information and requests for resources and reagents should be directed to and will be fulfilled by the Lead Contact, Chikashi Terao.

## METHOD DETAILS

### MENTR ML models

We designed the MENTR ML models by combining deep convolutional neural networks (from +/- 100-kb genome sequence around TSS to epigenetic features, using publicly available pre-trained DeepSEA Beluga models (Zhou and Troyanskaya, 2015; Zhou et al., 2018)) and binary classifier using gradient boosting trees (from epigenetic features to transcription probabilities) (Chen and Guestrin, 2016). In the DeepSEA Beluga model, 2,002 epigenetic features in 200-bp bin were predicted from 200-bp +/- 900-bp genome sequences (Zhou et al., 2018); then we obtained total 2,002,000 epigenetic features in +/- 100-kb regions (2,002 × 1,000 bins) using Pytorch (v0.4.0) using cuda 9.0. Based on the hypothesis that transcription would be affected by near epigenetic events, we aggregated each type of epigenetic features by using 5 types of exponential transformation depending bin–TSS distance (Zhou et al., 2018). In this transformation, we did not use transcript strand information because expression levels of transcribed enhancer had no strand information. Notably, using strand information rather slightly decreased predictive accuracies (Figure S11). Mathematical representation of the exponential transformation is as follows:

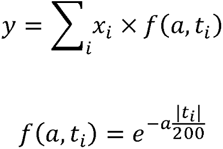

where *x_i_* is an epigenetic feature of bin *i*(*i* = 1,…, 1,000), *t_i_* is the mean distance between bin *i*, and TSS and *a* = {0.01, 0.02, 0.05, 0.10, 0.20| is an empirical parameter of 5 types of transformation (Zhou et al., 2018). The aggregated 10,010 epigenetic features (2,002 × 5) were used for input variables for gradient boosting trees (GBT) implemented in python xgboost library (v0.72.1). After training GBTs with the parameters (Table S13), we performed isotonic regression (Niculescu-Mizil and Caruana, 2012) using valid datasets (see below) by python sklearn (v0.19.1) to calibrate the output values from the GBT models (Figure S12). We considered that the calibration would be an important procedure to compare probabilities among different models and to use probability as a threshold for selecting reliable results of *in silico* mutagenesis. We used the calibrated probabilities for all analysis unless otherwise specified. We noted that MENTR_linear_ ML models with linear penalized logistic regression models using boosting (Bühlmann, 2006; Chen and Guestrin, 2016) were used for Figure S2 and Figure S8 to reduce computational costs in massive screening conditions. We measured the accuracies of MENTR ML models by AUROC by using R package pROC (v1.8).

### Datasets for training and testing MENTR ML models

We obtained representative TSS positions of promoters and inferred midpoint positions of enhancers from FANTOM5 phase 2.5 data files. We called both as TSS unless otherwise specified. We used the 1,829 samples from the major human primary cell-types and tissues and CAGE transcriptomes, and calculated expression levels of CAGE clusters as counts per million and normalized them by relative log expression methods across all the libraries (see the details in Hon et al., 2017). We calculated mean expression levels of a transcript for 347 types of sample ontologies, which is a set of non-redundant cell (n = 173) and tissue (n = 174). In the training MENTR ML models, we considered >0 expression levels as “on” and the others as “off”. We used 241,794 autosomal CAGE clusters on chromosome 8 as test data (for evaluate predictive accuracy), and 80% out of the others as train data (for training ML models) and the remaining as valid data (for early stopping and probability calibration). We used FANTOM CAT annotations (Hon et al., 2017) to define mRNA promoters, lncRNA promoters (antisense lncRNA, intergenic lncRNA, divergent lncRNA, and sense intronic lncRNA), and other types of promoters. In the category specific analysis, we excluded CAGE clusters with multiple annotations.

### Evaluation of FP predictions using nascent RNA profiling

We trained ML models using five ENCODE cell lines (HeLa, HepG2, MCF-7, K562, and GM12878) profiled by CAGE, and defined CAGE peaks whose expression level were 0 but predicted probabilities > 0.5 in test data as FP predictions. We developed two MENTR ML models from 2 replicates of CAGE transcriptome for each cell line and used FP predictions in either ML models in the following analysis. We evaluated the expression levels of FP predictions in expression levels of the nascent elongating RNAs from NET-CAGE samples treated by 2 M urea lysis buffer (Hirabayashi et al., 2019). We noted that the NET-CAGE could capture transcribed CAGE peaks before degradation and therefore detect lowly expressed transcripts including enhancers at the same sequence depth. We evaluated enrichment of expressed nascent RNAs in FP predictions by Fisher’s Exact Test using Bonferroni correction (P < (0.05/14)).

### *in silico* mutagenesis using the trained MENTR ML models

We predicted probability from hg19 sequence with the reference allele and the alternative allele using the trained MENTR ML models and calculated the difference (*in silico* mutation effect). In the prospective purpose like cataloging mutation effects of GWAS variants, we calculated in silico mutation effects on all possible transcripts whose TSS located within +/- 100-kb surrounding the variant.

### Verification of *in silico* mutagenesis using promoter and enhancer-level eQTL and caQTL results

We obtained published LCL CAGE transcriptomes for 154 unrelated European donors (Garieri et al., 2017) and quantified and normalized them as described in the above. We calculated mean expression levels among the LCL from 154 donors and used them for training and testing MENTR ML models (LCL MENTR model). Besides, we obtained published 5,376 promoter usage QTL (puQTL) and 110 enhancer activity QTL (eaQTL) results via personal communication (Garieri et al., 2017). We obtained 1KG variants in LD with the lead SNPs (1KGp3v5 EUR; r^2^ ≥ 0.7, window≤500kb) and performed *in silico* mutagenesis for the variants on paired transcripts (promoter or enhancer) using the LCL MENTR model for 7,063 variant-promoter pairs and 161 variant-enhancer pairs (±100kb variants surrounding each peak).

Similarly, we obtained published chromatin accessibility QTL (caQTL) results (Kumasaka et al., 2019). We used 297,308 lead caQTL-peak ATAC-seq pairs from 227,128 ATAC-seq peaks (only autosomal). We searched ≥50% overlapped regions between the ATAC-seq regions and FANTOM5 promoter or enhancer regions, resulting into 20,963 FANTOM5 permissive enhancer-ATAC-seq pairs and 89,199 FANTOM5 promoter-ATAC-seq peaks. We found variants in LD (1KGp3v5 EUR; r^2^ 0.7, window 500kb) and finally 86,724 variant-promoter pairs and 24,726 variant-enhancer pairs (±100kb variants surrounding each peak).

### Verification of *in silico* mutagenesis using gene-level eQTL results

We performed *in silico* mutagenesis of eQTL variants catalogued in GTEx v7 (MAF ≥ 100) using the paired tissues or cell-types in FANTOM5 sample ontologies (Table S5). In order to compare the GTEx gene-level mutation effects with MENTR promoter-level mutation effect, we filtered the promoter-level mutation effects by the robust or permissive threshold value, aggregated the promoter-level predicted probabilities from input sequences with reference or alternative allele based on the FANTOM CAT annotations, and calculated the difference between them (Figure S8).

### Making catalog of GWAS-traits-associated transcripts

We downloaded GWAS associations and their ancestry information from GWAS catalog (r2019-07-12). From the downloaded 143,963 records, we excluded records including variants-interaction (epistasis) results, unknown risk allele, and variant whose P-value ≥ 5×10^-8^; records without rsID (dbSNP 151); record, and records not SNP or InDels. Finally, 53,186 records remained (autosomal+chrX; SNP+InDel). By using broad ancestral category information, we split the dataset into only EAS study (1,426 variants), only EUR study (35,823), and others (15,765). For only EAS or EUR study, we obtained variants in LD (r^2^ ≥ 0.7, window≤500kb) using 1KGp3 datasets for each population. Similarly, we obtained Biobank Japan GWAS variants including lead variants after conditioning analysis from the published papers (Akiyama et al., 2017; Ishigaki et al., 2019; Kanai et al., 2018) and variants in LD using EAS 1KGp3 datasets.

In the regression analysis in Figure 4, we used all the results obtained in the above *in silico* mutagenesis, calculated maximum absolute mutation effects (*max effects*) over the FANTOM5 347 sample ontologies and LCL, and converted the probability-scale values to logit values (*logit max effects*). We estimated regression coefficients of the logit max effects on GWAS effect size for the indicated complex diseases only using variants ≥1% in each testing dataset. The coefficients were conditioned on MAF of variants, absolute variant-TSS distance, and dummy variable for promoter or enhancer. All the estimates were shown in Table S10.

### *in silico mutagenesis* of variants in credible sets from IBD fine-mapping

We obtained 95% credible sets in 139 independent associated regions in IBD from the fine-mapping paper (Huang et al., 2017). We calculated *in silico* mutation effects of the biallelic variants in dbSNP 151. We reported the number of credible regions including variant–transcript pairs with at least permissive mutation effects (≥0.05) in Figure 4B, and showed all results of the pairs with robust mutation effects (≥0.1) in Table S11. For the transcript type-specific count in Figure 4C, we only used CAGE peaks whose corresponding transcript pairs were only the indicated transcript type.

## QUANTIFICATION AND STATISTICAL ANALYSIS

We used he statistical computing language R (https://www.r-project.org/) in all the statistical testing by the indicated statistical method.

## DATA AND CODE AVAILABILITY

The pre-trained MENTR ML models (347 sample ontologies and LCL) and the source code for training MENTR ML models and running *in silico* mutagenesis are publicly hosted at https://github.com/koido/XXX (*available after acceptance*). GWAS trait-associated ncRNA database is publicly available in a user-friendly GUI application (*available after acceptance*). These are publicly released under GPL v3 and are free for use for academic and non-commercial applications.

## Supplemental Information titles and legends

**Figure S1. Details about MENTR ML.**

(A) Comparisons between required datasets for MENTR ML models and conventional eQTL study. In MENTR ML models, CAGE transcriptome data is only required. Existing large-scale CAGE transcriptome, such as FANTOM5 datasets, can be used. In eQTL study, transcriptome data for the tissue and genotypes from the same individuals are required for estimating mutation effects for each transcript (β_eQTL_) in a tissue. (B) Workflow of MENTR ML training and evaluation. See the details in METHOD DETAILS section.

**Figure S2. Accurate prediction of ncRNA expression by combining MENTR ML models with CAGE transcriptome.**

(A) Prediction accuracies of ExPecto methods (Zhou et al., 2018) on GTEx RNA sequence datasets (re-analysis of predictive accuracies among 218 types of tissues) and FANTOM5 CAGE transcriptome datasets (347 sample ontologies). (B,C) Prediction accuracies of the indicated methods (x-axis) on lncRNAs (B) and mRNAs (C) in the FANTOM5 CAGE transcriptome datasets. Spearman’s ρ values were compared by violin plot and the mean values were shown by dot. P-values were calculated by Wilcoxon signed rank test.

**Figure S3. Maximized predictive accuracy by using ±100-kb sequence.**

We compared effects of input genome sequence length (x-axis for each plot), transcript type for training (each violin plot) on predicting promoter- (blue) and enhancer-level expression (red) in FANTOM5 347 sample ontologies. In these analyses, MENTR_linear_ was used. AUROC values were compared by violin plot and the mean values were shown by dot. P-values were calculated by Wilcoxon signed rank test.

**Figure S4. Partial dependency of predictive accuracy on annotation of target promoters.**

We compared the dependency of predictive accuracy on the annotation of CAGE transcript. We selected CAGE peaks with each of the indicated annotation and calculated AUROC of them.

**Figure S5. Illustrative overview of methods to evaluate accuracies of *in silico* mutation effects.**

(A) Comparison between *β_mutgen_* and *β_QTL_* from LCL CAGE QTL study (Garieri et al., 2017). (B) Comparison between *β_mutgen_* and *β_QTL_* from LCL caQTL study (Kumasaka et al., 2019). See the details in METHOD DETAILS section.

**Figure S6. Verification of *in silico* mutation effects on all promoters.**

Concordance rate (y-axis) of directions of *in silico* mutation effect (β_*mutgen*_) for all promoters and effect size from the QTL study (*β_eQTL_*) at the threshold of absolute *β_mutgen_*. *β_QTL_* was obtained from LCL CAGE QTL study (Garieri et al., 2017) in (A) and from LCL caQTL study (Kumasaka et al., 2019) in (B). See Figure 3.

**Figure S7. Correlation between *in silico* mutation effect size and caQTL effect size.**

*β_mutgen_* (x-axis) was compared with *β_QTL_* from LCL caQTL study (Kumasaka et al., 2019), for enhancers in (A) and promoters in (B) In these plots, we assume that *β_QTL_* of variants in LD. ρ. Spearman’s ρ. red dot, absolute *β_mutgen_* ≥ 0.1; triangle dot, absolute variant-TSS distance > 1kb. We excluded variants with 0 mutation effect from this analysis.

**Figure S8. Gene-level verification of MENTR *in silico* mutation effects for various types of tissues.**

(A) Workflow of calculating gene-level mutation effects (*Δy*). *Δy* values were calculated from promoter-level mutation effects (*Δy_p_*) after filtered by the baseline, permissive, and robust threshold. (B) Concordance rate (y-axis) of directions of the *Δy* and effect size of eQTL from GTEx v7 at the indicated threshold of absolute *y*. The concordance rates of 26 tissues (Table S5) were shown by violin plot and the mean values were shown as dot. P-values were calculated by Wilcoxon signed rank test.

**Figure S9. MENTR predictions of causal variants which were experimentally validated.**

Association plots (*upper panel*) and absolute mutation effects (*lowered panel*) for *CCR1* (A) and *HLA-DOA* (B). The functional variant (rs147398495 and rs381218) and variants in LD (r^2^ ≥ 0.2 in 1KGp3v5 EUR) were shown. The permissive threshold of *in silico* mutation effect (0.5) was shown as a dashed line. Larger dots mean that the mutation effect sizes are greater than the robust threshold (0.1). The strongest *in silico* mutation effects among 348 models for each mutation were shown.

**Figure 10. MENTR predictions of causal variants in Crohn’s disease by linking known functional variants with transcribed enhancers in relevant cell-types.**

*in silico* mutation effects of rs4456788, rs713875, and their variants in LD. For the association plots, summary statistics of Crohn’s disease transethnic meta GWAS from (Liu et al., 2015) were used in (A; for rs4456788) and those from (Franke et al., 2010) were used in (B; for rs713875). eQTL for near-by *ICOSLG* and *LIF* were obtained from GTEx v7 eQTL significant results. *in silico* mutation effects for the indicated transcripts were predicted from the MENTR ML model for CD4^+^, α-β - T cell in (A) and CD8^+^, - Tα-β cell in (B), both of which were the cell type with the strongest *in silico* mutation effects (Table S11). See also Figure 5.

**Figure S11. Decreased predictive accuracies in MENTR ML models by using transcript strand information.**

We compared effects of considering strand in our MENTR ML models (x-axis, AUROC from ML models using strand information like ExPecto models (Zhou et al., 2018); y-axis, our methods which do not use strand information). We trained ML models by using only promoters. In this figure, MENTR_linear_ was used.

**Figure S12. Calibration of probabilities for accurate metrics.**

Representative calibration plots were shown. Percentage of expressed CAGE peaks for each bin were shown in y-axis, and mean values of probabilities for each bin (evenly spaced by 0.2 of range [0, 1]) were shown in x-axis. The number of CAGE peaks in each bin were shown in the upper bar plots. GBT: only gradient boosting trees; GBT+IR: isotonic regression after gradient boosting trees, used for MENTR ML models. Sample ontology ID for each plot was UBERON:0010133 in (A), UBERON:0009722 in (B), and UBERON:0005911 in (C).

## References

1. (DGT), T.F.C. and the R.P. and C., Forrest, A.R.R., Kawaji, H., Rehli, M., Baillie, J.K., Hoon, M.J.L. de, Haberle, V., Lassmann, T., Kulakovskiy, I. V., Lizio, M., et al. (2014). A promoter-level mammalian expression atlas. Nature 507, 462–470.

2. Aguet, F., Brown, A.A., Castel, S.E., Davis, J.R., He, Y., Jo, B., Mohammadi, P., Park, Y., Parsana, P., Segrè, A. V., et al. (2017). Genetic effects on gene expression across human tissues. Nature 550, 204–213.

3. Akiyama, M., Okada, Y., Kanai, M., Takahashi, A., Momozawa, Y., Ikeda, M., Iwata, N., Ikegawa, S., Hirata, M., Matsuda, K., et al. (2017). Genome-wide association study identifies 112 new loci for body mass index in the Japanese population. Nat. Genet. 49, 1458–1467.

4. Alasoo, K., Rodrigues, J., Mukhopadhyay, S., Knights, A.J., Mann, A.L., Kundu, K., Hale, C., Dougan, G., and Gaffney, D.J. (2018). Shared genetic effects on chromatin and gene expression indicate a role for enhancer priming in immune response. Nat. Genet. 50, 424–431.

5. Andersson, R., Gebhard, C., Miguel-Escalada, I., Hoof, I., Bornholdt, J., Boyd, M., Chen, Y., Zhao, X., Schmidl, C., Suzuki, T., et al. (2014). An atlas of active enhancers across human cell types and tissues. Nature 507, 455–461.

6. Anish, R., Hossain, M.B., Jacobson, R.H., and Takada, S. (2009). Characterization of Transcription from TATA-Less Promoters: Identification of a New Core Promoter Element XCPE2 and Analysis of Factor Requirements. PLoS One 4, e5103.

7. Ardlie, K.G., Deluca, D.S., Segre, A. V, Sullivan, T.J., Young, T.R., Gelfand, E.T., Trowbridge, C.A., Maller, J.B., Tukiainen, T., Lek, M., et al. (2015). The Genotype-Tissue Expression (GTEx) pilot analysis: Multitissue gene regulation in humans. Science (80-.). 348, 648–660.

8. van Arensbergen, J., Pagie, L., FitzPatrick, V.D., de Haas, M., Baltissen, M.P., Comoglio, F., van der Weide, R.H., Teunissen, H., Võsa, U., Franke, L., et al. (2019). High-throughput identification of human SNPs affecting regulatory element activity. Nat. Genet. 51, 1160–1169.

9. Banovich, N.E., Li, Y.I., Raj, A., Ward, M.C., Greenside, P., Calderon, D., Tung, P.Y., Burnett, J.E., Myrthil, M., Thomas, S.M., et al. (2018). Impact of regulatory variation across human iPSCs and differentiated cells. Genome Res. 28, 122–131.

10. Boyd, M., Thodberg, M., Vitezic, M., Bornholdt, J., Vitting-Seerup, K., Chen, Y., Coskun, M., Li, Y., Lo, B.Z.S., Klausen, P., et al. (2018). Characterization of the enhancer and promoter landscape of inflammatory bowel disease from human colon biopsies. Nat. Commun. 9.

11. Bryois, J., Garrett, M.E., Song, L., Safi, A., Giusti-Rodriguez, P., Johnson, G.D., Shieh, A.W., Buil, A., Fullard, J.F., Roussos, P., et al. (2018). Evaluation of chromatin accessibility in prefrontal cortex of individuals with schizophrenia. Nat. Commun. 9, 3121.

12. Bühlmann, P. (2006). Boosting for high-dimensional linear models. Ann. Stat. 34, 559–583.

13. Chen, T., and Guestrin, C. (2016). XGBoost: A scalable tree boosting system. In Proceedings of the ACM SIGKDD International Conference on Knowledge Discovery and Data Mining (New York, New York, USA: Association for Computing Machinery), pp. 785–794.

14. Chen, L., Ge, B., Casale, F.P., Vasquez, L., Kwan, T., Garrido-Martín, D., Watt, S., Yan, Y., Kundu, K., Ecker, S., et al. (2016). Genetic Drivers of Epigenetic and Transcriptional Variation in Human Immune Cells. Cell 167, 1398–1414.e24.

15. Degner, J.F., Pai, A.A., Pique-Regi, R., Veyrieras, J.-B., Gaffney, D.J., Pickrell, J.K., De Leon, S., Michelini, K., Lewellen, N., Crawford, G.E., et al. (2012). DNase-QTLs are a major determinant of human expression variation. Nature 482, 390–394.

16. Delaneau, O., Zazhytska, M., Borel, C., Giannuzzi, G., Rey, G., Howald, C., Kumar, S., Ongen, H., Popadin, K., Marbach, D., et al. (2019). Chromatin three-dimensional interactions mediate genetic effects on gene expression. Science 364, eaat8266.

17. Demenais, F., Margaritte-Jeannin, P., Barnes, K.C., Cookson, W.O.C., Altmüller, J., Ang, W., Barr, R.G., Beaty, T.H., Becker, A.B., Beilby, J., et al. (2018). Multiancestry association study identifies new asthma risk loci that colocalize with immune-cell enhancer marks. Nat. Genet. 50, 42–50.

18. Donczew, R., and Hahn, S. (2017). Mechanistic Differences in Transcription Initiation at TATA-Less and TATA-Containing Promoters. Mol. Cell. Biol. 38.

19. Fahy, J. V. (2009). Eosinophilic and neutrophilic inflammation in asthma insights from clinical studies. In Proceedings of the American Thoracic Society, pp. 256–259.

20. Farh, K.K.H., Marson, A., Zhu, J., Kleinewietfeld, M., Housley, W.J., Beik, S., Shoresh, N., Whitton, H., Ryan, R.J.H., Shishkin, A.A., et al. (2015). Genetic and epigenetic fine mapping of causal autoimmune disease variants. Nature 518, 337–343.

21. Finucane, H.K., Bulik-Sullivan, B., Gusev, A., Trynka, G., Reshef, Y., Loh, P.-R., Anttila, V., Xu, H., Zang, C., Farh, K., et al. (2015). Partitioning heritability by functional annotation using genome-wide association summary statistics. Nat. Genet. 47, 1228–1235.

22. Finucane, H.K., Reshef, Y.A., Anttila, V., Slowikowski, K., Gusev, A., Byrnes, A., Gazal, S., Loh, P.-R., Lareau, C., Shoresh, N., et al. (2018). Heritability enrichment of specifically expressed genes identifies disease-relevant tissues and cell types. Nat. Genet. 50, 621–629.

23. Franke, A., McGovern, D.P.B., Barrett, J.C., Wang, K., Radford-Smith, G.L., Ahmad, T., Lees, C.W., Balschun, T., Lee, J., Roberts, R., et al. (2010). Genome-wide meta-analysis increases to 71 the number of confirmed Crohn’s disease susceptibility loci. Nat. Genet. 42, 1118–1125.

24. Garieri, M., Delaneau, O., Santoni, F., Fish, R.J., Mull, D., Carninci, P., Dermitzakis, E.T., Antonarakis, S.E., and Fort, A. (2017). The effect of genetic variation on promoter usage and enhancer activity. Nat. Commun. 8, 1358.

25. Gate, R.E., Cheng, C.S., Aiden, A.P., Siba, A., Tabaka, M., Lituiev, D., Machol, I., Gordon, M.G., Subramaniam, M., Shamim, M., et al. (2018). Genetic determinants of co-accessible chromatin regions in activated T cells across humans. Nat. Genet. 50, 1140–1150.

26. Hirabayashi, S., Bhagat, S., Matsuki, Y., Takegami, Y., Uehata, T., Kanemaru, A., Itoh, M., Shirakawa, K., Takaori-Kondo, A., Takeuchi, O., et al. (2019). NET-CAGE characterizes the dynamics and topology of human transcribed cis-regulatory elements. Nat. Genet. 51, 1369–1379.

27. Hoffman, G.E., Bendl, J., Girdhar, K., Schadt, E.E., and Roussos, P. (2019). Functional interpretation of genetic variants using deep learning predicts impact on chromatin accessibility and histone modification. Nucleic Acids Res. 47, 10597–10611.

28. Hon, C.C., Ramilowski, J.A., Harshbarger, J., Bertin, N., Rackham, O.J.L., Gough, J., Denisenko, E., Schmeier, S., Poulsen, T.M., Severin, J., et al. (2017). An atlas of human long non-coding RNAs with accurate 5′ ends. Nature 543, 199–204.

29. Huang, H., Fang, M., Jostins, L., Umićvić Mirkov, M., Boucher, G., Anderson, C.A., Andersen, V., Cleynen, I., Cortes, A., Crins, F., et al. (2017). Fine-mapping inflammatory bowel disease loci to single-variant resolution. Nature 547, 173–178.

30. Hutchinson, J.N., Raj, T., Fagerness, J., Stahl, E., Viloria, F.T., Gimelbrant, A., Seddon, J., Daly, M., Chess, A., and Plenge, R. (2014). Allele-Specific Methylation Occurs at Genetic Variants Associated with Complex Disease. PLoS One 9, e98464.

31. Imam, T., Park, S., Kaplan, M.H., and Olson, M.R. (2018). Effector T helper cell subsets in inflammatory bowel diseases. Front. Immunol. 9, 1212.

32. Iotchkova, V., Ritchie, G.R.S., Geihs, M., Morganella, S., Min, J.L., Walter, K., Timpson, N.J., Dunham, I., Birney, E., and Soranzo, N. (2019). GARFIELD classifies disease-relevant genomic features through integration of functional annotations with association signals. Nat. Genet. 51, 343–353.

33. Ishigaki, K., Kochi, Y., Suzuki, A., Tsuchida, Y., Tsuchiya, H., Sumitomo, S., Yamaguchi, K., Nagafuchi, Y., Nakachi, S., Kato, R., et al. (2017). Polygenic burdens on cell-specific pathways underlie the risk of rheumatoid arthritis. Nat. Genet. 49, 1120–1125.

34. Ishigaki, K., Akiyama, M., Kanai, M., Takahashi, A., Kawakami, E., Sugishita, H., Sakaue, S., Matoba, N., Low, S.-K., Okada, Y., et al. (2019). Large scale genome-wide association study in a Japanese population identified 45 novel susceptibility loci for 22 diseases. BioRxiv 795948.

35. Kadivar, M., Petersson, J., Svensson, L., and Marsal, J. (2016). CD8αβ + γδ T Cells: A Novel T Cell Subset with a Potential Role in Inflammatory Bowel Disease. J. Immunol. 197, 4584–4592.

36. Kanai, M., Akiyama, M., Takahashi, A., Matoba, N., Momozawa, Y., Ikeda, M., Iwata, N., Ikegawa, S., Hirata, M., Matsuda, K., et al. (2018). Genetic analysis of quantitative traits in the Japanese population links cell types to complex human diseases. Nat. Genet. 50, 390–400.

37. Kelley, D.R., Reshef, Y.A., Bileschi, M., Belanger, D., McLean, C.Y., and Snoek, J. (2018). Sequential regulatory activity prediction across chromosomes with convolutional neural networks. Genome Res. 28, 739–750.

38. Kim, T.-K., Hemberg, M., Gray, J.M., Costa, A.M., Bear, D.M., Wu, J., Harmin, D.A., Laptewicz, M., Barbara-Haley, K., Kuersten, S., et al. (2010). Widespread transcription at neuronal activity-regulated enhancers. Nature 465, 182–187.

39. Kristjánsdóttir, K., Kwak, Y., Tippens, N.D., Lis, J.T., Kang, H.M., and Kwak, H. (2018). Population-scale study of eRNA transcription reveals bipartite functional enhancer architecture. BioRxiv 426908.

40. Kumasaka, N., Knights, A.J., and Gaffney, D.J. (2019). High-resolution genetic mapping of putative causal interactions between regions of open chromatin. Nat. Genet. 51, 128–137.

41. Lamparter, D., Marbach, D., Rueedi, R., Kutalik, Z., and Bergmann, S. (2016). Fast and Rigorous Computation of Gene and Pathway Scores from SNP-Based Summary Statistics. PLoS Comput. Biol. 12, 1–20.

42. Liu, J.Z., Van Sommeren, S., Huang, H., Ng, S.C., Alberts, R., Takahashi, A., Ripke, S., Lee, J.C., Jostins, L., Shah, T., et al. (2015). Association analyses identify 38 susceptibility loci for inflammatory bowel disease and highlight shared genetic risk across populations. Nat. Genet. 47, 979–986.

43. Mattioli, K., Volders, P.-J., Gerhardinger, C., Lee, J.C., Maass, P.G., Melé, M., and Rinn, J.L. (2019). High-throughput functional analysis of lncRNA core promoters elucidates rules governing tissue specificity. Genome Res. 29, 344–355.

44. Maurano, M.T., Humbert, R., Rynes, E., Thurman, R.E., Haugen, E., Wang, H., Reynolds, A.P., Sandstrom, R., Qu, H., Brody, J., et al. (2012). Systematic Localization of Common Disease-Associated Variation in Regulatory DNA. Science (80-.). 337, 1190–1195.

45. Miller, C.L., Pjanic, M., Wang, T., Nguyen, T., Cohain, A., Lee, J.D., Perisic, L., Hedin, U., Kundu, R.K., Majmudar, D., et al. (2016). Integrative functional genomics identifies regulatory mechanisms at coronary artery disease loci. Nat. Commun. 7.

46. Murakawa, Y., Yoshihara, M., Kawaji, H., Nishikawa, M., Zayed, H., Suzuki, H., FANTOM Consortium, and Hayashizaki, Y. (2016). Enhanced Identification of Transcriptional Enhancers Provides Mechanistic Insights into Diseases. Trends Genet. 32, 76–88.

47. Niculescu-Mizil, A., and Caruana, R.A. (2012). Obtaining Calibrated Probabilities from Boosting.

48. Pelikan, R.C., Kelly, J.A., Fu, Y., Lareau, C.A., Tessneer, K.L., Wiley, G.B., Wiley, M.M., Glenn, S.B., Harley, J.B., Guthridge, J.M., et al. (2018). Enhancer histone-QTLs are enriched on autoimmune risk haplotypes and influence gene expression within chromatin networks. Nat. Commun. 9, 2905.

49. Saunders, G., Baudis, M., Becker, R., Beltran, S., Béroud, C., Birney, E., Brooksbank, C., Brunak, S., Van den Bulcke, M., Drysdale, R., et al. (2019). Leveraging European infrastructures to access 1 million human genomes by 2022. Nat. Rev. Genet. 20, 693–701.

50. Terao, C., Yoshifuji, H., Nakajima, T., Yukawa, N., Matsuda, F., and Mimori, T. (2016). Ustekinumab as a therapeutic option for Takayasu arteritis: from genetic findings to clinical application. Scand. J. Rheumatol. 45, 80–82.

51. Turner, A.W., Martinuk, A., Silva, A., Lau, P., Nikpay, M., Eriksson, P., Folkersen, L., Perisic, L., Hedin, U., Soubeyrand, S., et al. (2016). Functional analysis of a novel genome-wide association study signal in SMAD3 that confers protection from coronary artery disease. Arterioscler. Thromb. Vasc. Biol. 36, 972–983.

52. Wéra, O., Lancellotti, P., and Oury, C. (2016). The Dual Role of Neutrophils in Inflammatory Bowel Diseases. J. Clin. Med. 5, 118.

53. Yadav, P.K., Chen, C., and Liu, Z. (2011). Potential role of NK cells in the pathogenesis of inflammatory bowel disease. J. Biomed. Biotechnol. 2011, 348530.

54. Zhang, Z., Lee, J.H., Ruan, H., Ye, Y., Krakowiak, J., Hu, Q., Xiang, Y., Gong, J., Zhou, B., Wang, L., et al. (2019). Transcriptional landscape and clinical utility of enhancer RNAs for eRNA-targeted therapy in cancer. Nat. Commun. 10.

55. Zhou, J., and Troyanskaya, O.G. (2015). Predicting effects of noncoding variants with deep learning-based sequence model. Nat. Methods 12, 931–934.

56. Zhou, J., Theesfeld, C.L., Yao, K., Chen, K.M., Wong, A.K., and Troyanskaya, O.G. (2018). Deep learning sequence-based ab initio prediction of variant effects on expression and disease risk. Nat. Genet. 1.

