## Supplementary material for "Predicting cell-type-specific non-coding RNA transcription from genome sequence": Key Resources Table

| REAGENT or RESOURCE | SOURCE | IDENTIFIER |
|---|---|---|
| Deposited Data | | |
| LCL CAGE transcriptomes | (Garieri et al., 2017) | E-MTAB-5835 |
| GWAS catalog | (Buniello et al., 2019) | r2019-07-12 |
| CAGE transcriptome in FANTOM5 | ((DGT) et al., 2014; Andersson et al., 2014; Hon et al., 2017) | https://fantom.gsc.riken.jp/5/datafiles |
| Representative TSS of CAGE peaks (promoters) | | https://fantom.gsc.riken.jp/5/datafiles/phase2.5/extra/CAGE_peaks/hg19.cage_peak_phase1and2combined_coord.bed.gz |
| Inferred mid position of enhancers | | https://fantom.gsc.riken.jp/5/datafiles/phase2.5/extra/Enhancers/human_permissive_enhancers_phase_1_and_2.bed.gz |
| CAGE peak annotations | | https://fantom.gsc.riken.jp/5/datafiles/phase2.5/extra/CAGE_peaks_annotation |
| caQTL from LCL ATAC-seq | (Kumasaka et al., 2019) | N/A |
| eQTL GTEx v7 | (Aguet et al., 2017) | https://www.gtexportal.org/home/ |
| CAGE and NET-CAGE transcriptome of the five ENCODE cell lines (GM12878, HeLa-S3, HepG2, K562, and MCF-7) | (Hirabayashi et al., 2019) | GSE118075 |

| | | |
|---|---|---|
| DeepSEA (Beluga) | (Zhou and Troyanskaya, 2015; Zhou et al., 2018) | https://github.com/FunctionLab/ExPecto |
| IBD transethnic meta GWAS results | (Franke et al., 2010; Liu et al., 2015) | https://www.ibdgenetics.org/downloads.html |
| Software and Algorithms | | |
| MENTR software | This study | GitHub (available after acceptance) |
| Pretrained MENTR models | This study | GitHub (available after acceptance) |
| Other | | |
| 5,376 puQTL and 110 eaQTL from LCL CAGE | (Garieri et al., 2017) | N/A |
