## Supplementary Figures for "Predicting cell-type-specific non-coding RNA transcription from genome sequence"

### Figure S1.

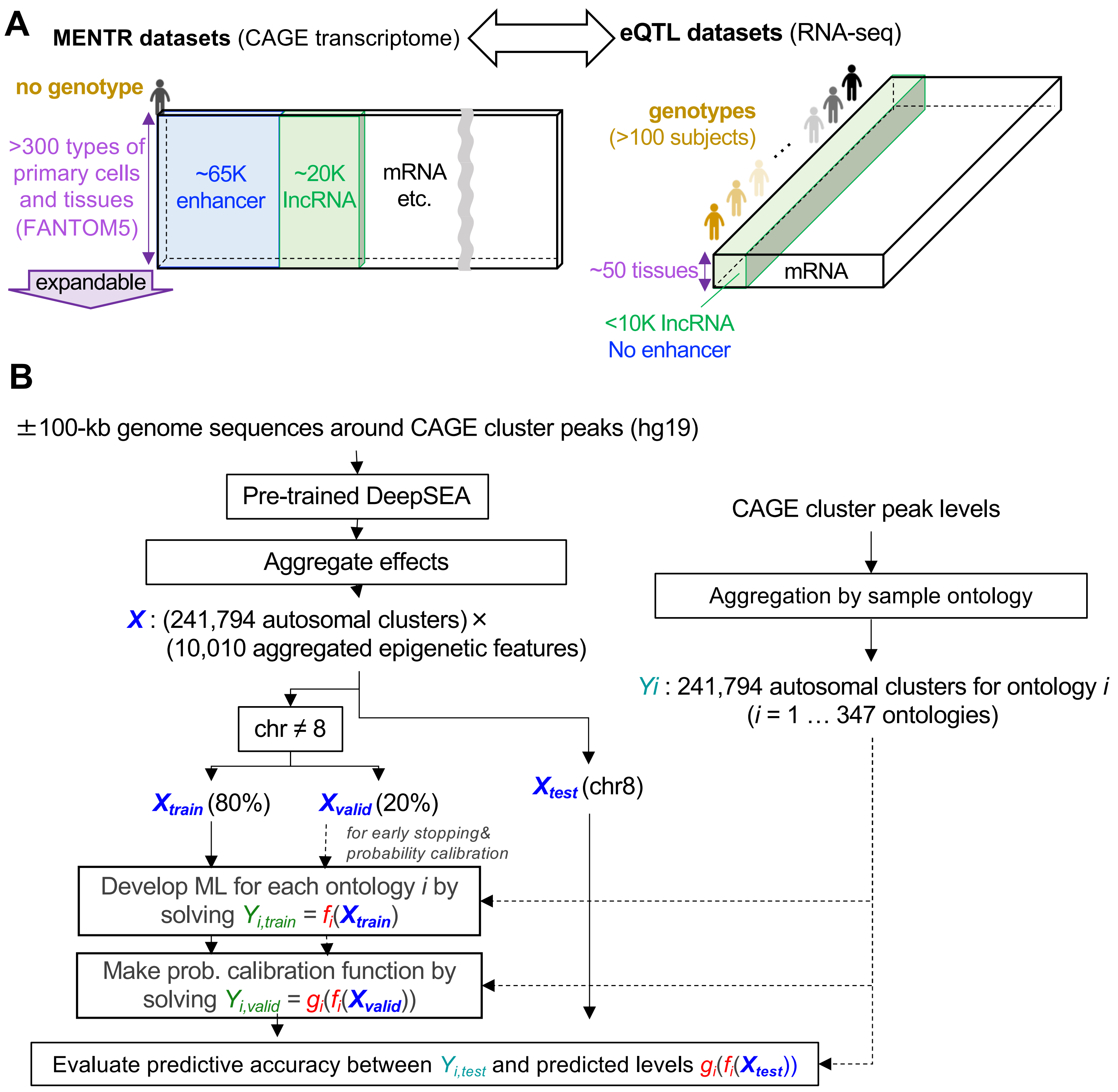
A
MENTR datasets (CAGE transcriptome)
eQTL datasets (RNA-seq)
no genotype
>300 types of primary cells and tissues (FANTOM5)
expandable
~65K enhancer
~20K lncRNA
mRNA etc.
genotypes (>100 subjects)
~50 tissues
mRNA
<10K lncRNA
No enhancer
B
±100-kb genome sequences around CAGE cluster peaks (hg19)
Pre-trained DeepSEA
Aggregate effects
X : (241,794 autosomal clusters) × (10,010 aggregated epigenetic features)
CAGE cluster peak levels
Aggregation by sample ontology
Yi : 241,794 autosomal clusters for ontology i (i = 1 … 347 ontologies)
chr ≠ 8
Xtrain (80%)
Xvalid (20%)
Xtest (chr8)
for early stopping& probability calibration
Develop ML for each ontology i by solving Yi,train = fi(Xtrain)
Make prob. calibration function by solving Yi,valid = gi(fi(Xvalid))
Evaluate predictive accuracy between Yi,test and predicted levels gi(fi(Xtest))

#### Figure S2.

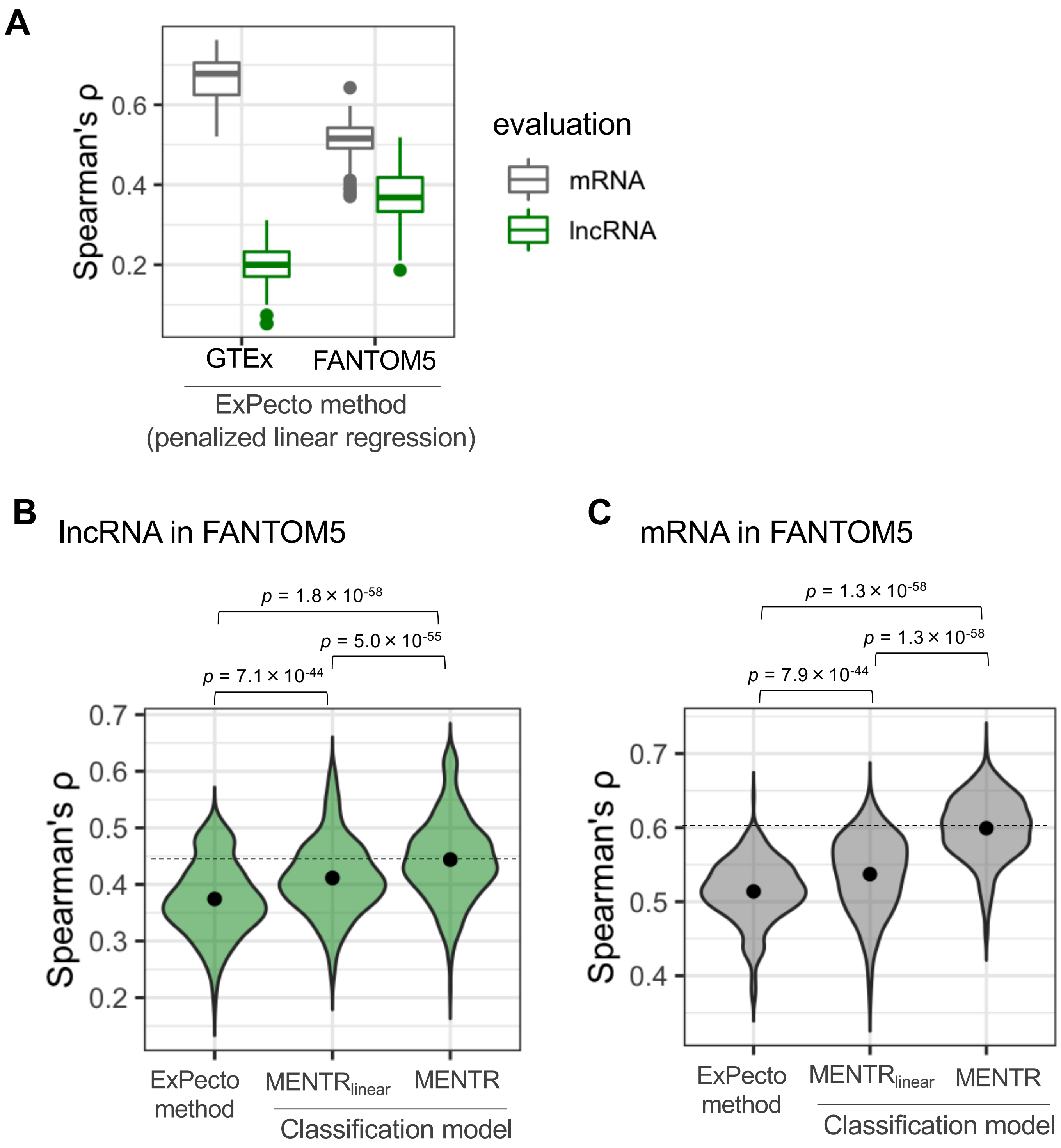
A
Spearman's ρ
0.6
0.4
0.2
GTEx
FANTOM5
ExPecto method
(penalized linear regression)
evaluation
mRNA
lncRNA
B
lncRNA in FANTOM5
p = 1.8 × 10-58
p = 5.0 × 10-55
p = 7.1 × 10-44
Spearman's ρ
0.7
0.6
0.5
0.4
0.3
0.2
ExPecto method
MENTRlinear
MENTR
Classification model
C
mRNA in FANTOM5
p = 1.3 × 10-58
p = 1.3 × 10-58
p = 7.9 × 10-44
Spearman's ρ
0.7
0.6
0.5
0.4
ExPecto method
MENTRlinear
MENTR
Classification model

### Figure S3.

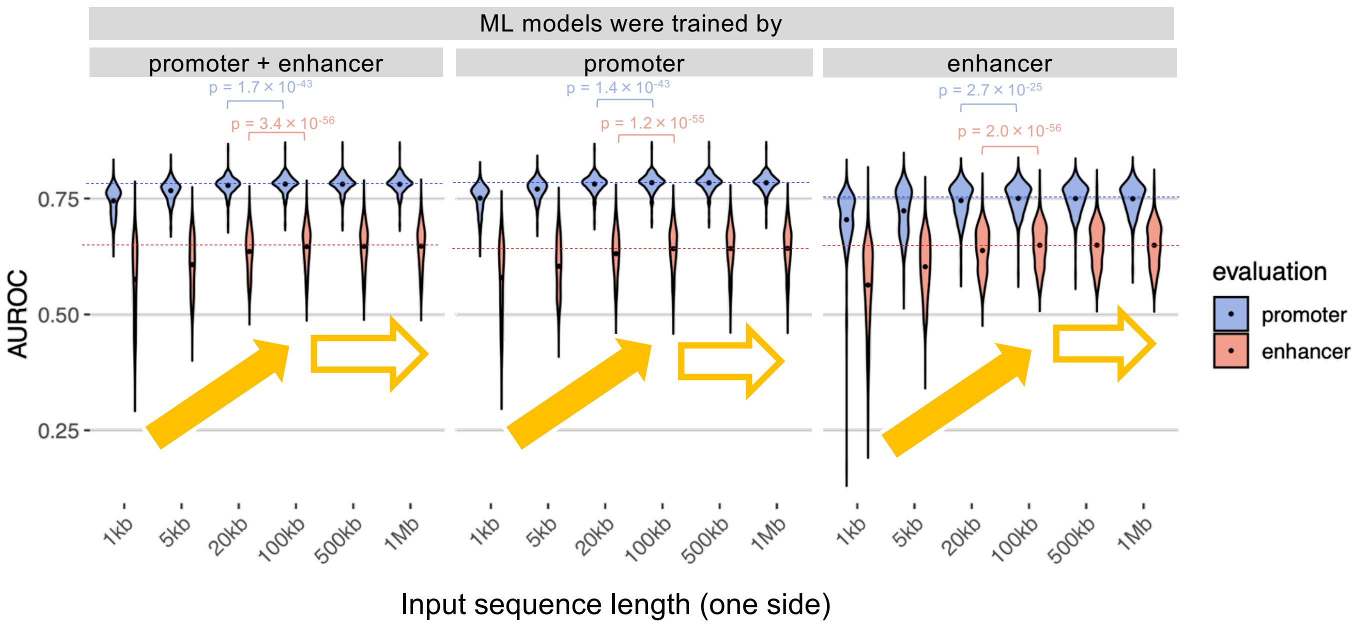
ML models were trained by
promoter + enhancer
promoter
enhancer
p = 1.7 × 10-43
p = 3.4 × 10-56
p = 1.4 × 10-43
p = 1.2 × 10-55
p = 2.7 × 10-25
p = 2.0 × 10-56
AUROC
0.75
0.50
0.25
1kb
5kb
20kb
100kb
500kb
1Mb
Input sequence length (one side)
evaluation
promoter
enhancer

### Figure S4.

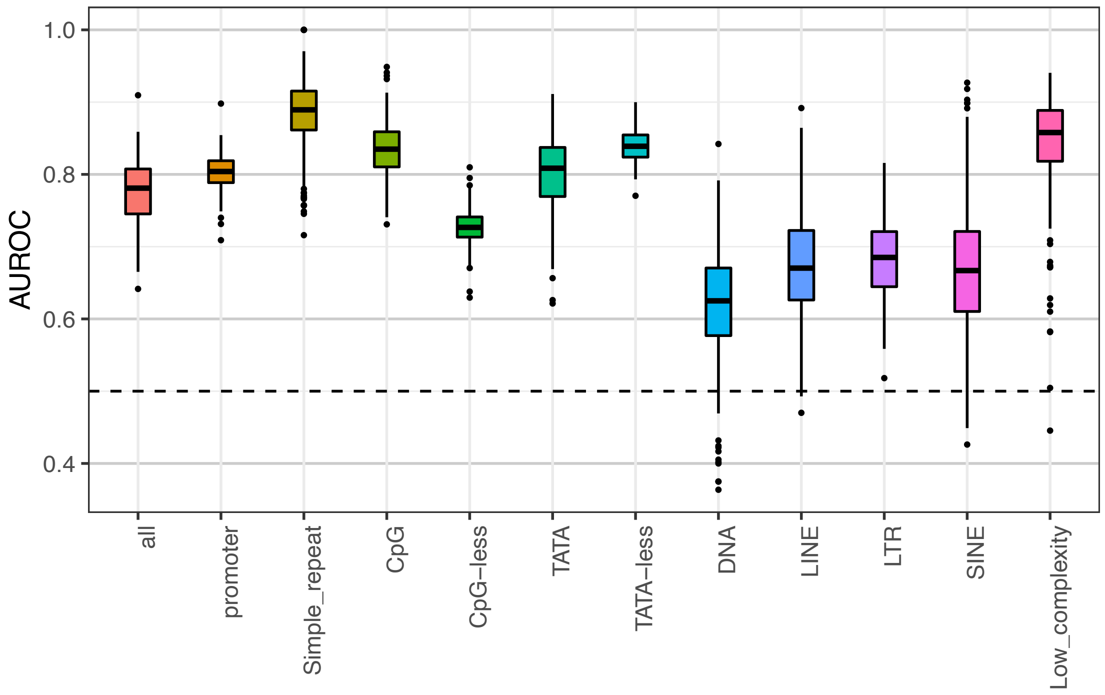
1.0
0.8
0.6
0.4
AUROC
all
promoter
Simple_repeat
CpG
CpG−less
TATA
TATA−less
DNA
LINE
LTR
SINE
Low_complexity

### Figure S5.

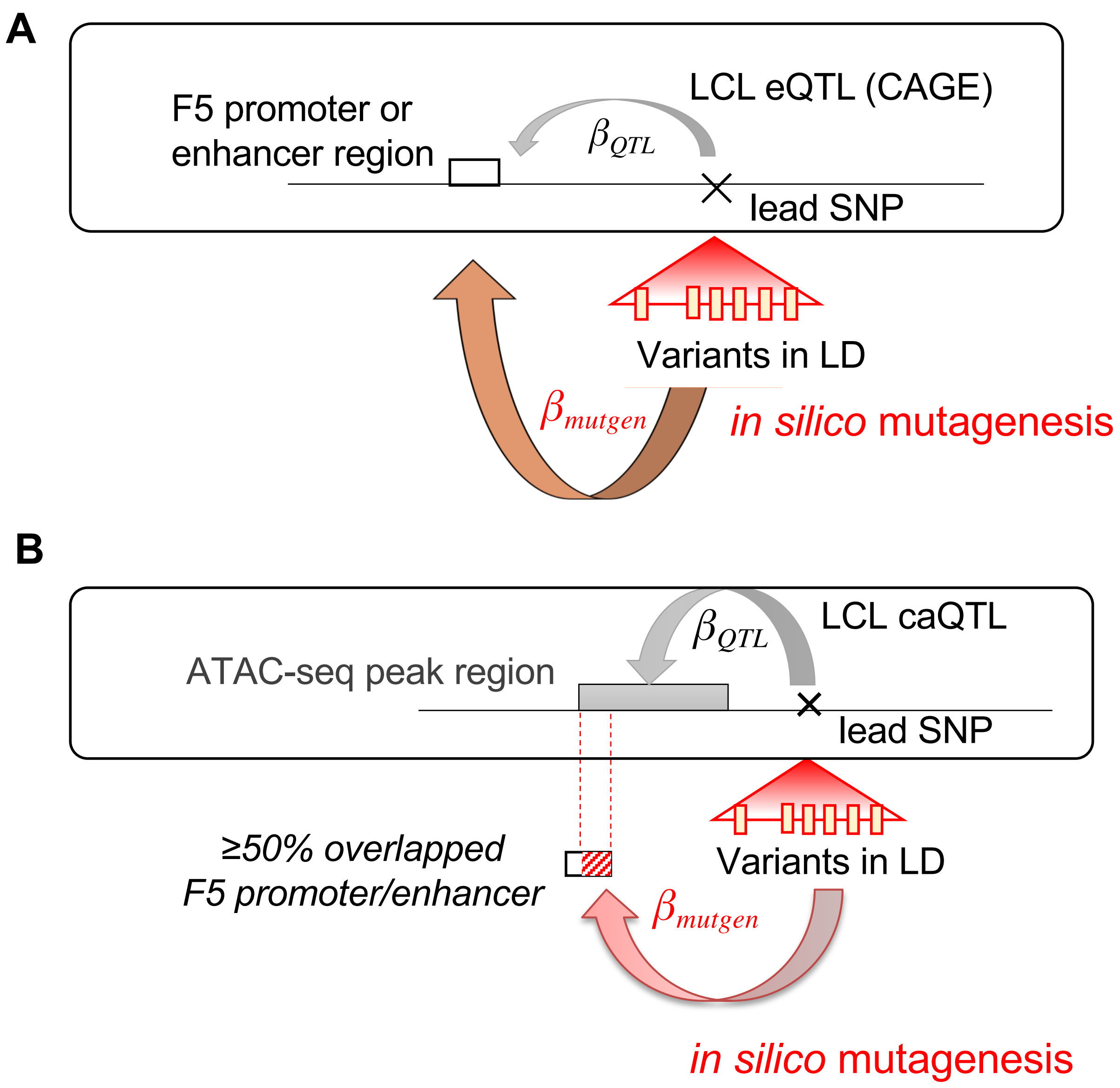
A
F5 promoter or
enhancer region
LCL eQTL (CAGE)
$\beta_{QTL}$
lead SNP
Variants in LD
$\beta_{mutgen}$
*in silico* mutagenesis
B
ATAC-seq peak region
$\beta_{QTL}$
LCL caQTL
lead SNP
≥50% overlapped
F5 promoter/enhancer
Variants in LD
$\beta_{mutgen}$
*in silico* mutagenesis

**Figure S6.**

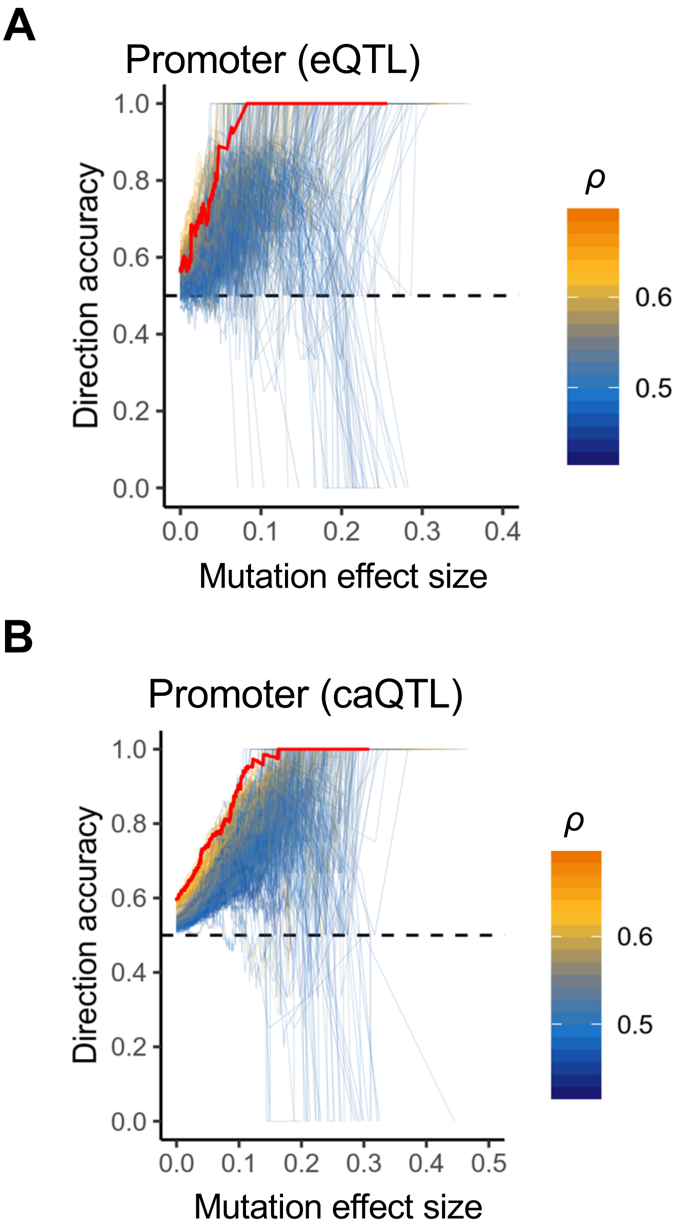
A
Promoter (eQTL)
Direction accuracy
1.0
0.8
0.6
0.4
0.2
0.0
0.0
0.1
0.2
0.3
0.4
Mutation effect size
ρ
0.6
0.5
B
Promoter (caQTL)
Direction accuracy
1.0
0.8
0.6
0.4
0.2
0.0
0.0
0.1
0.2
0.3
0.4
0.5
Mutation effect size
ρ
0.6
0.5

**Figure S7.**

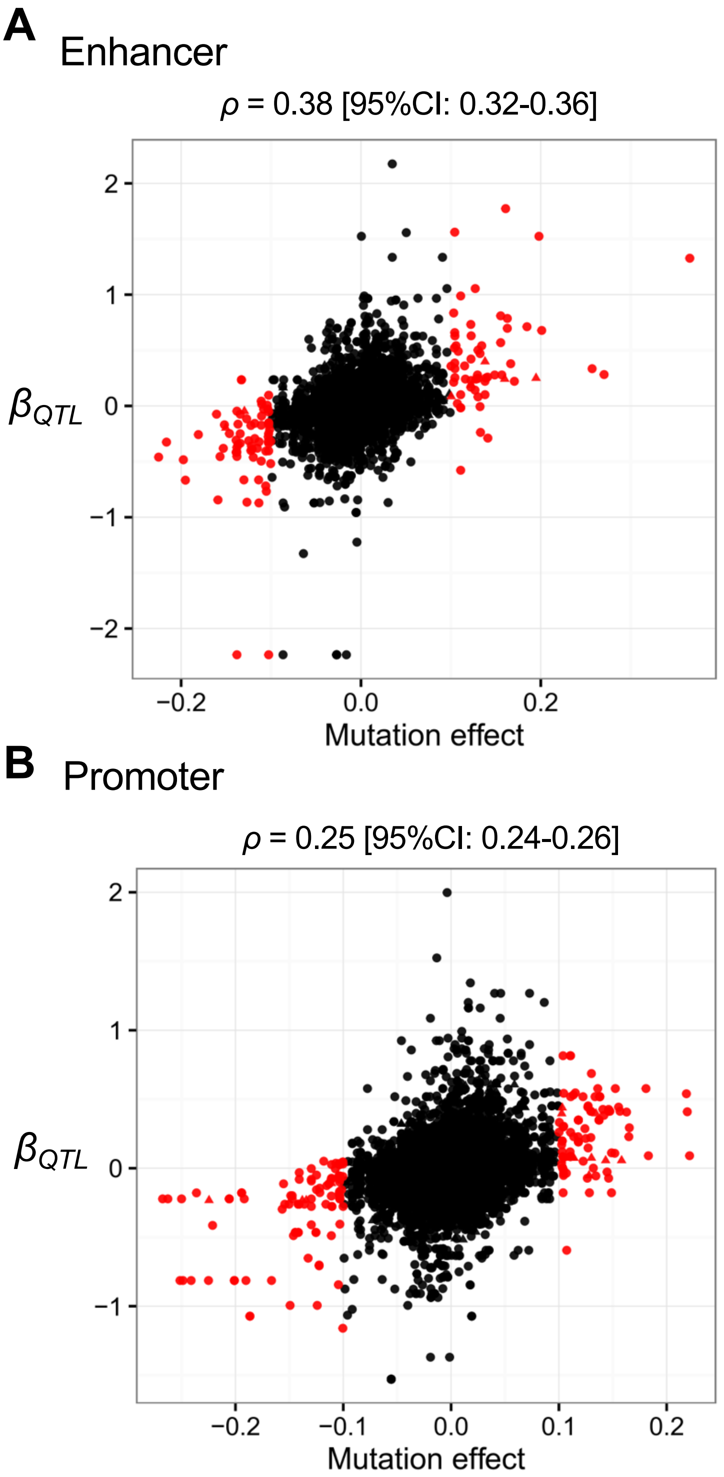
A
Enhancer
ρ = 0.38 [95%CI: 0.32-0.36]
2
1
0
−1
−2
βQTL
−0.2
0.0
0.2
Mutation effect
B
Promoter
ρ = 0.25 [95%CI: 0.24-0.26]
2
1
0
−1
βQTL
−0.2
−0.1
0.0
0.1
0.2
Mutation effect

### Figure S8.

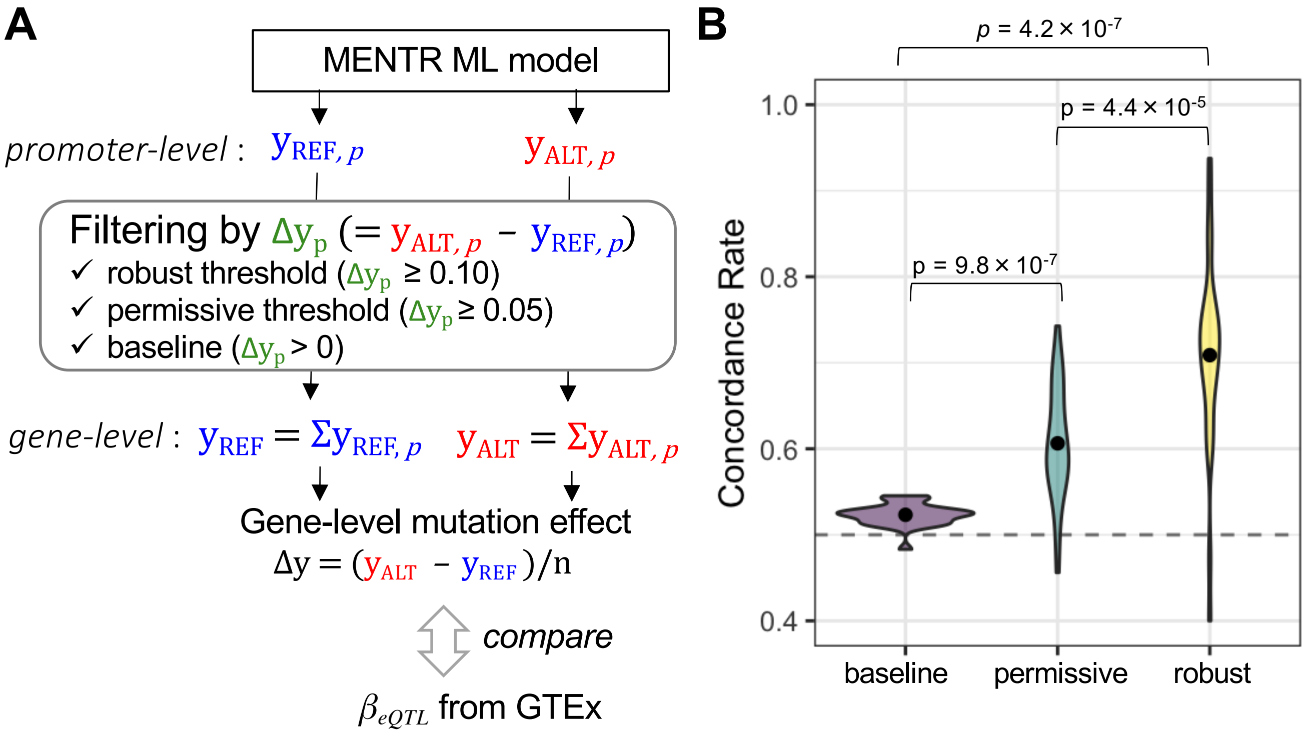

A
MENTR ML model
promoter-level : $y_{REF,p}$ $y_{ALT,p}$
Filtering by $\Delta y_p$ (= $y_{ALT,p}$ − $y_{REF,p}$)
✓ robust threshold ($\Delta y_p \geq 0.10$)
✓ permissive threshold ($\Delta y_p \geq 0.05$)
✓ baseline ($\Delta y_p > 0$)
gene-level : $y_{REF} = \Sigma y_{REF,p}$ $y_{ALT} = \Sigma y_{ALT,p}$
Gene-level mutation effect
$\Delta y = (y_{ALT} - y_{REF})/n$
compare
$\beta_{eQTL}$ from GTEx
B
$p = 4.2 \times 10^{-7}$
$p = 4.4 \times 10^{-5}$
$p = 9.8 \times 10^{-7}$
Concordance Rate
1.0
0.8
0.6
0.4
baseline
permissive
robust

### Figure S9.

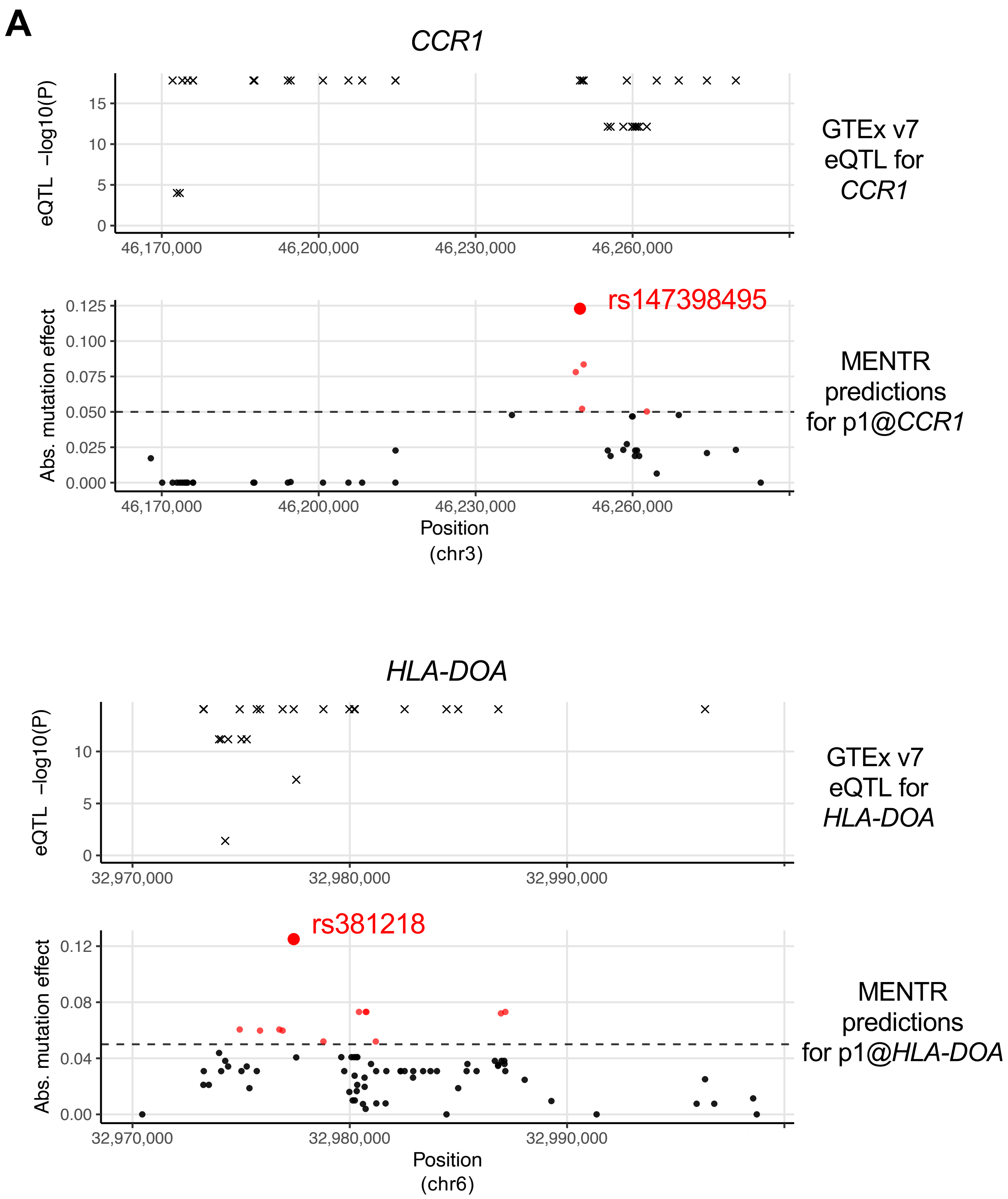
A
CCR1
eQTL −log10(P)
0
5
10
15
46,170,000
46,200,000
46,230,000
46,260,000
GTEx v7 eQTL for CCR1
Abs. mutation effect
0.000
0.025
0.050
0.075
0.100
0.125
rs147398495
MENTR predictions for p1@CCR1
Position (chr3)
B
HLA-DOA
eQTL −log10(P)
0
5
10
32,970,000
32,980,000
32,990,000
GTEx v7 eQTL for HLA-DOA
Abs. mutation effect
0.00
0.04
0.08
0.12
rs381218
MENTR predictions for p1@HLA-DOA
Position (chr6)

### Figure S10.

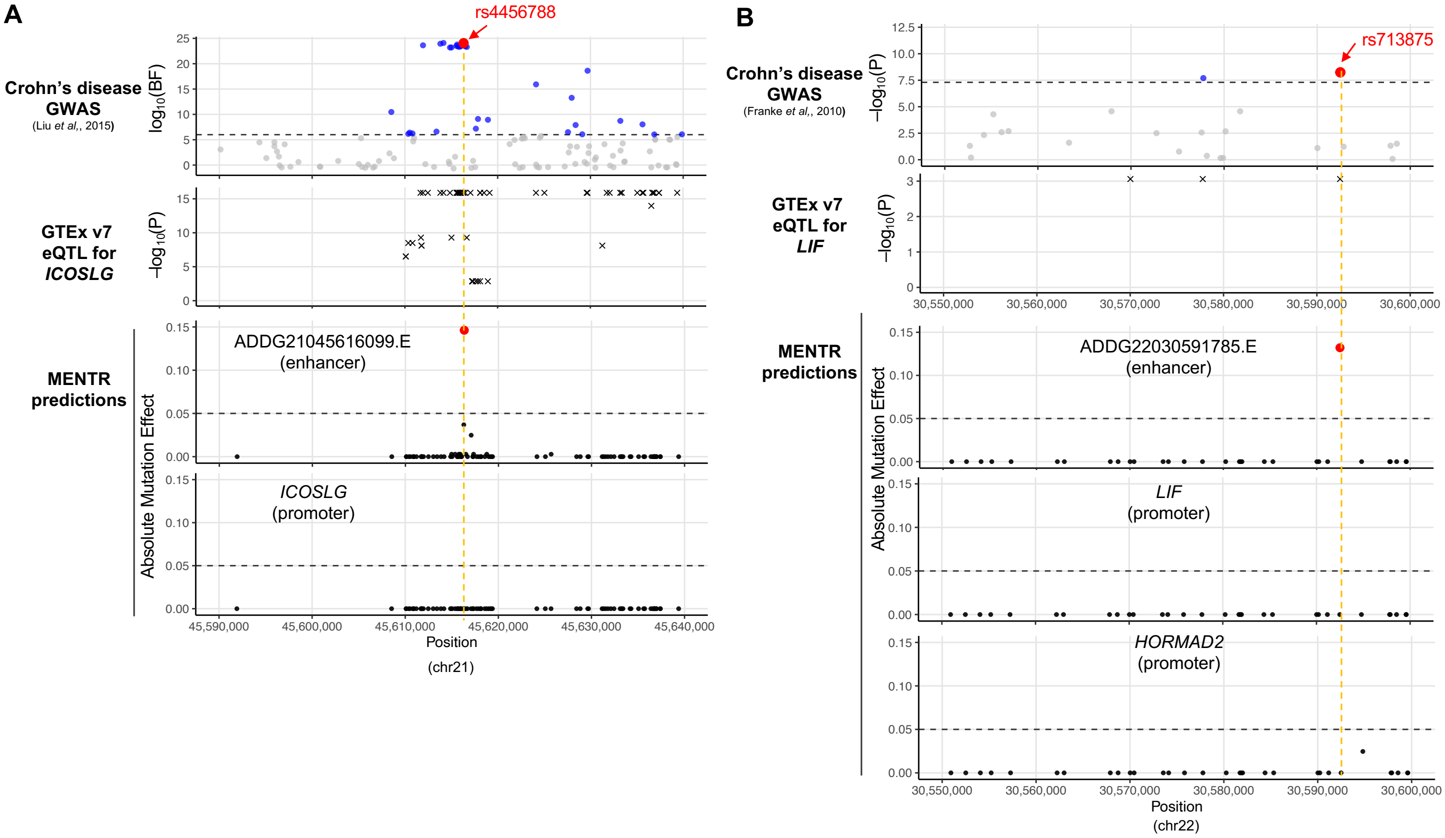
A
rs4456788
Crohn's disease GWAS
(Liu et al., 2015)
log10(BF)
GTEx v7 eQTL for ICOSLG
−log10(P)
MENTR predictions
Absolute Mutation Effect
ADDG21045616099.E
(enhancer)
ICOSLG
(promoter)
45,590,000
45,600,000
45,610,000
45,620,000
45,630,000
45,640,000
Position
(chr21)
B
rs713875
Crohn's disease GWAS
(Franke et al., 2010)
−log10(P)
GTEx v7 eQTL for LIF
ADDG22030591785.E
(enhancer)
LIF
(promoter)
HORMAD2
(promoter)
30,550,000
30,560,000
30,570,000
30,580,000
30,590,000
30,600,000
Position
(chr22)

### Figure S11.

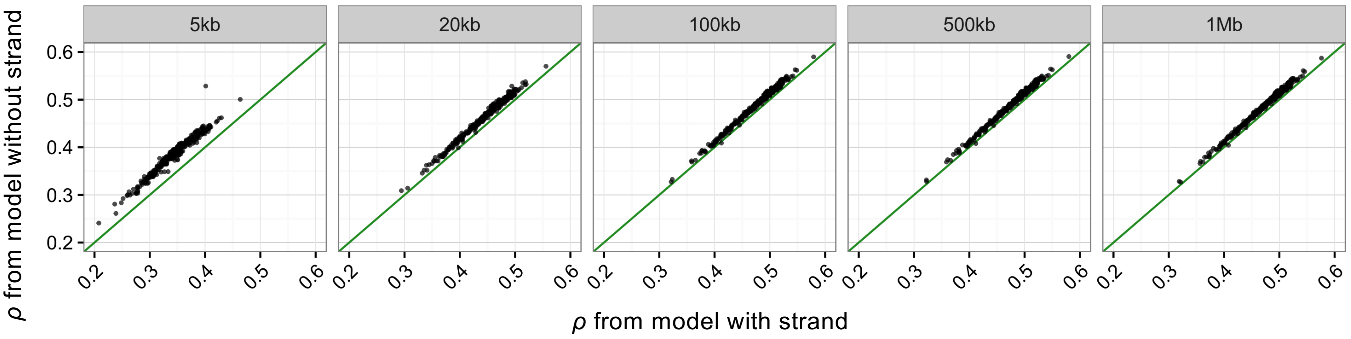
5kb
20kb
100kb
500kb
1Mb
ρ from model without strand
ρ from model with strand
0.2
0.3
0.4
0.5
0.6

### Figure S12.

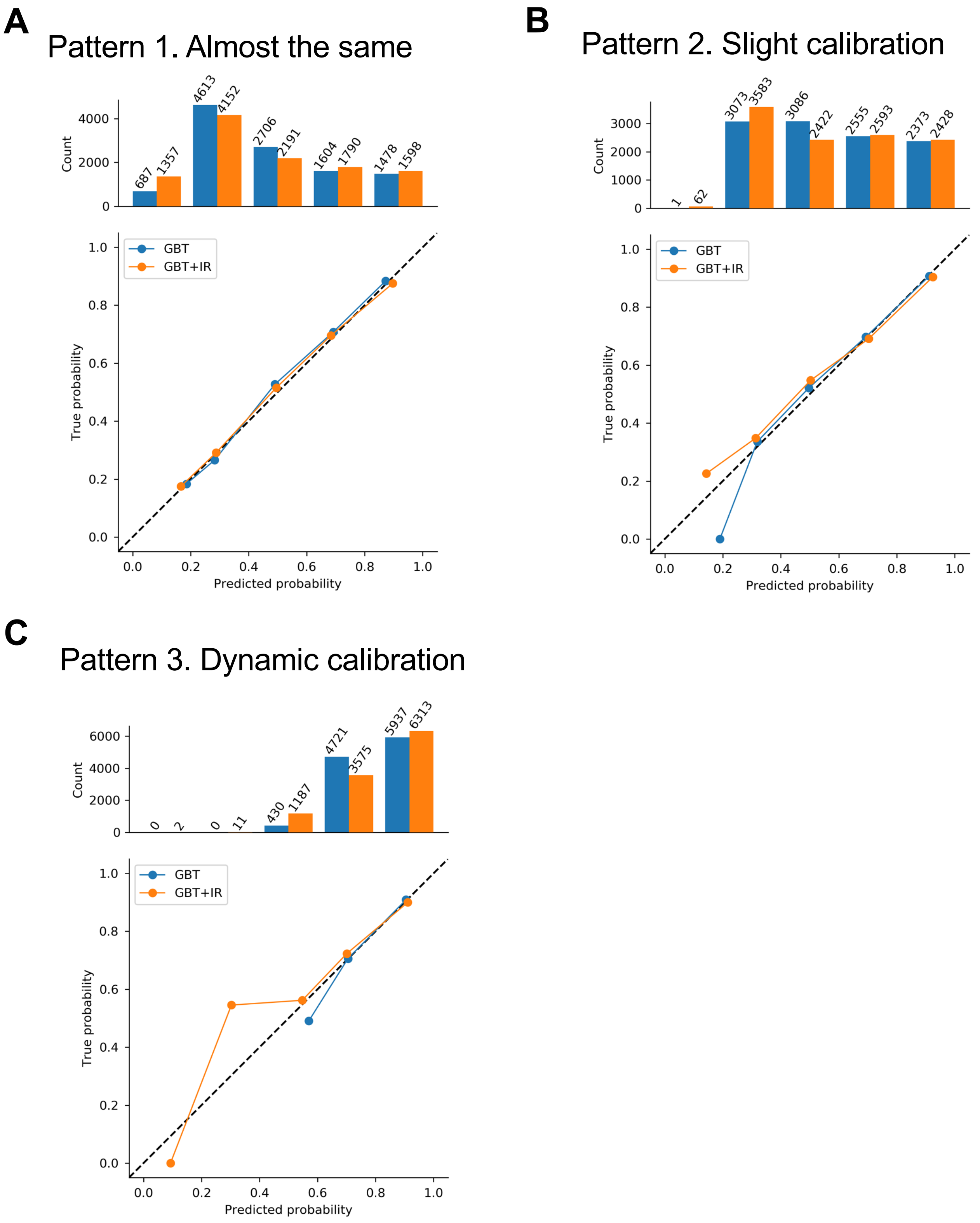
A
Pattern 1. Almost the same
Count
0
2000
4000
687
1357
4613
4152
2706
2191
1604
1790
1478
1598
GBT
GBT+IR
True probability
Predicted probability
0.0
0.2
0.4
0.6
0.8
1.0
B
Pattern 2. Slight calibration
Count
0
1000
2000
3000
1
62
3073
3583
3086
2422
2555
2593
2373
2428
GBT
GBT+IR
True probability
Predicted probability
C
Pattern 3. Dynamic calibration
Count
0
2000
4000
6000
0
2
0
11
430
1187
4721
3575
5937
6313
GBT
GBT+IR
True probability
Predicted probability
